# THEMIS is a priming substrate of non-receptor tyrosine phosphatase PTPN6/SHP1 and plays dual roles during T cell development

**DOI:** 10.1101/2021.12.21.473566

**Authors:** Jiali Zhang, Erwei Zuo, Minfang Song, Li Chen, Zhenzhou Jiang, Shengmiao Chen, Xuexue Xiong, Yuetong Wang, Piliang Hao, Tiffany Horng, Min Zhuang, Liye Zhang, Haopeng Wang, Gaofeng Fan

**Author notes:** Correspondence: Gaofeng Fan, School of Life Science and Technology, ShanghaiTech University, Shanghai, China 201210. 86-13917147596. Haopeng Wang, School of Life Science and Technology, ShanghaiTech University, Shanghai, China 201210. 86-15001913510. Equal Contribution.

## Abstract

THEMIS plays an indispensable role in T cells, but its mechanism of action is highly controversial. Using the systematic proximity labeling methodology PEPSI, we identified THEMIS as an uncharacterized substrate for the phosphatase SHP1. Saturated mutagenesis analysis revealed that THEMIS phosphorylation at the evolutionally conserved Tyr34 residue was oppositely regulated by SHP1 and the kinase LCK. Like THEMIS^-/-^ mice, THEMIS^Y34F/Y34F^ knock-in mice showed a significant decrease in CD4 thymocytes and mature CD4 T cells, but a normal thymic development and peripheral homeostasis of CD8 T cells. Mechanistically, phosphorylated THEMIS induced by TCR activation acts as a “priming substrate” to bind SHP1 and convert its phosphatase activity from basal level to nearly fully activated level, ensuring an appropriate negative regulation of TCR signaling. However, cytokine signaling in CD8 T cells failed to elicit THEMIS Y34 phosphorylation, revealing both phosphorylation-dependent and -independent roles of THEMIS in controlling T cell maturation and expansion.

## Introduction

T cell development is a critical event in adaptive immunity, deficiency of which will trigger many autoimmune and immunodeficiency diseases. Thymocytes will mainly go through three stages characterized by expression status of the co-receptors CD4 and CD8: CD4^-^ CD8^-^(double negative (DN)) stage, CD4^+^CD8^+^ (double-positive (DP)) stage, and CD4^+^CD8^-^ or CD4^-^CD8^+^ (single-positive (SP)) stage. The dynamic change of the cell-surface signaling receptors, characterized by pre- and mature forms of the T cell antigen receptor (TCR), coordinates the driving force for the progression of thymocytes through these stages of maturation (Godfrey et al., 1993; Zuniga-Pflucker, 2004).

After successful rearrangement and expression of a TCR β-chain (TCRβ) in conjunction with upregulation of the coreceptor CD4 and CD8, immature DN T cells transit into DP cells, followed by a selection process which is key to T cell maturation. The outcome of this selection is largely dependent on the duration and strength of TCR signaling, which is influenced by the affinity of the expressed TCR for self-peptide ligands bound to major histocompatibility complex (self-pMHC) presented by thymic stromal cells. Thymocytes whose TCR binds too tightly to self-pMHC initiate activation-induced apoptosis known as “negative selection”, whereas those cells that receive too weak binding signals die by neglect. Only cells expressing TCR with intermediate affinity to self-pMHC are “positively selected” and progress to the SP stage (Germain, 2002). Besides TCR, the cytokine interleukin-7 (IL-7) mediated pro-survival pathway is also required in the later phase of CD8SP thymocytes maturation (Singer et al., 2008). Multiple signaling modulators downstream of TCR, including the protein tyrosine kinases (PTKs) LCK and ZAP-70, and the protein tyrosine phosphatase (PTP) PTPN6/SHP1, play indispensable roles in controlling the intensity and duration of the TCR signaling transduction and therefore T cell selection. Recently, THEMIS has been identified by several teams as a signaling hub important in the maturation of T cells (Allen, 2009; Fu et al., 2009; Johnson et al., 2009; Kakugawa et al., 2009; Lesourne et al., 2009; Patrick et al., 2009). Genetic ablation of THEMIS severely reduces the number of mature CD4SP thymocytes and, to a lesser extent, CD8SP thymocytes and peripheral T cells.

Reversible tyrosine phosphorylation, orchestrated by PTKs and PTPs, is critical for cells to respond to stimuli in their environment, such as TCR-pMHC engagement during thymocyte selection. For the past 40 years, we have witnessed a substantial breakthrough in PTK research, from substrate identification to functional characterization, and culminating in therapeutic potential with the development of pharmacological inhibitors. However, the importance and physiological significance of PTP remain under-appreciated. Actually, one debate regarding the function of THEMIS in T cell development during positive selection is focused on its role in regulating the phosphatase activity of SHP1. Although several groups agree on the essentiality of THEMIS during T cell development through its interaction with SHP1, two contradictory models have been proposed regarding the positive or negative role of THEMIS in moderating the phosphatase activity of SHP1, which leads to a decrease or increase of TCR signaling response accordingly (Choi et al., 2017a; Choi et al., 2017b; Fu et al., 2013; Gascoigne and Acuto, 2015; Gascoigne et al., 2016a; Mehta et al., 2018; Paster et al., 2015). Thus, there is an urgent need to develop a systematic platform to profile the whole spectrum of potential substrates for any given PTP and to further characterize the molecular function of these PTPs within a physiological context.

By combining strategies of substrate trapping for PTPs and pupylation-based interaction tagging (PUP-IT) for protein-protein interaction, we developed a systematic proximity labeling methodology PEPSI (PUP-IT Enhanced Protein Tyrosine Phosphatase Substrate Identification) to identify novel substrates for PTPs. We applied this approach to SHP1 and, surprisingly, identified THEMIS as an uncharacterized substrate for the phosphatase. Further studies demonstrated phosphorylation of Tyr 34 of THEMIS was oppositely controlled by LCK and SHP1 during T cell activation. To address the importance of THEMIS Y34 phosphorylation, we generated a THEMIS mutant mouse with tyrosine 34 mutated to phenylalanine. Similar to THEMIS^-/-^ mice, phosphorylation-deficient THEMIS^Y34F/Y34F^ mice showed a slightly milder but significant decrease in CD4 thymocytes and mature CD4 T cells, nevertheless had a normal CD8 T cell development and peripheral CD8 T cell homeostasis. Mechanistically, tyrosine phosphorylated THEMIS served as a “priming substrate” that binds SHP1 and activates SHP1’s function downstream of TCR signaling. By contrast, cytokine signaling that mediating CD8 T cells’ development and hemostasis didn’t induce THEMIS Y34 phosphorylation.

## Results

### PEPSI strategy identified THEMIS as a novel substrate of non-receptor tyrosine phosphatase SHP1

Pioneer work from the Tonks lab has demonstrated that the conserved Asp residue within the WPD loop of protein tyrosine phosphatases is important for substrate dissociation from the phosphatase after dephosphorylation (Flint et al., 1997). Mutating this residue to Ala within the enzyme will substantially prolong the binding and create a trap for potential substrates. This substrate trapping strategy helps differentiate the substrates from the binders of PTPs. Pupylation-based interaction tagging (PUP-IT) system, first invented by the Zhuang lab, is a powerful approach to identify transient and weak interactions like those between many enzymes and their substrates (Liu et al., 2018). A biotin-labeled small protein tag, Pup, is tagged to proteins that interact with a PafA-fused bait. Subsequent avidin bead enrichment and mass spectrometry detection can then identify candidate protein(s) associated with the protein of interest. Combining both substrate trapping strategy and PUP-IT approach enabled us to develop a systematic proximity labeling methodology to identify novel substrates for any given protein tyrosine phosphatase, which we named <u>P</u>UP-IT <u>E</u>nhanced Protein Tyrosine <u>P</u>hosphatase <u>S</u>ubstrate Identification (PEPSI) (Figure 1A).

**Figure 1.**
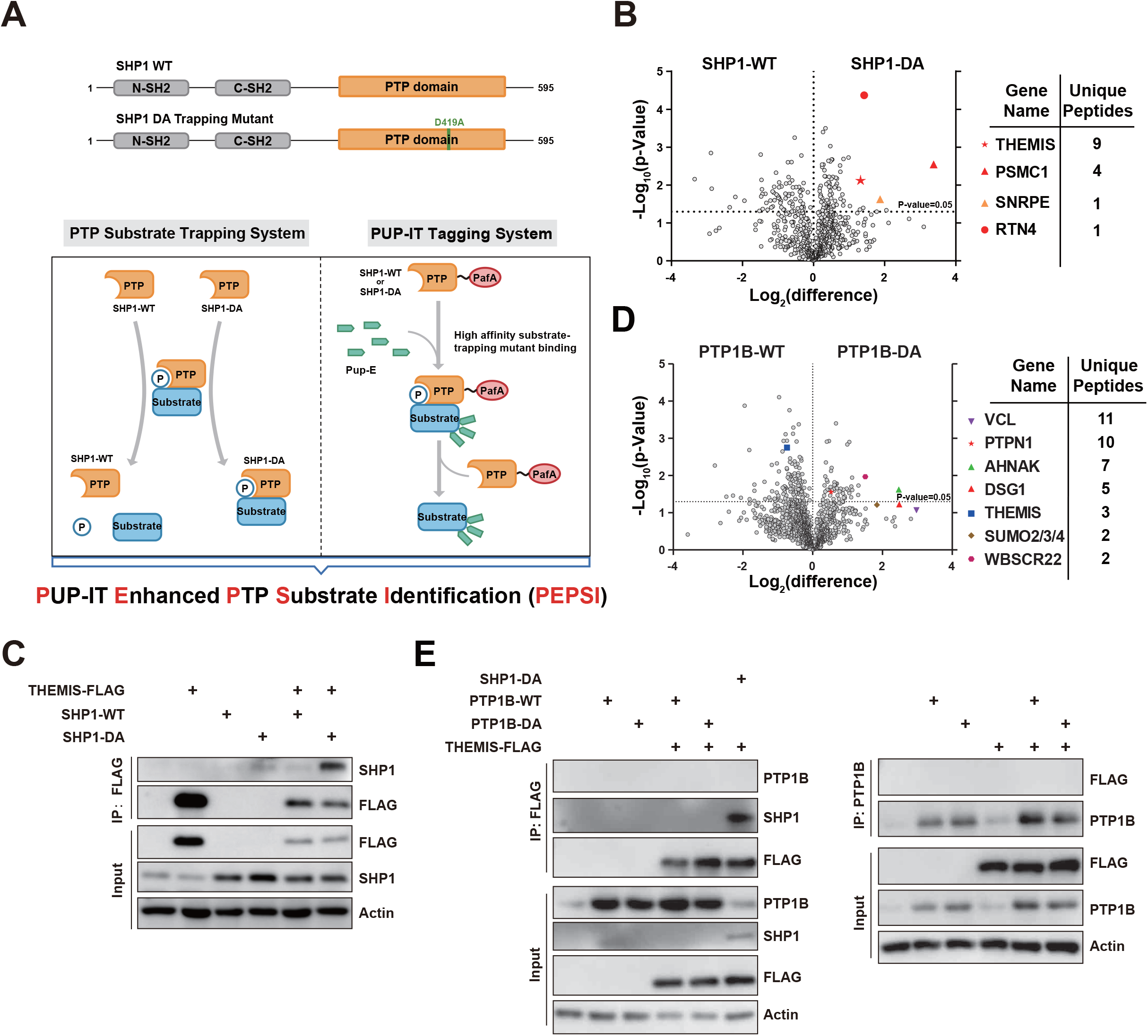
PEPSI strategy identified THEMIS as a novel substrate of non-receptor tyrosine phosphatase SHP1. (A) Schematic of the overall principle for the PEPSI method to protein tyrosine phosphatase SHP1-WT vs SHP1-D419A. The upper panel shows the domain distribution of SHP1 and the mutation site of the “trapping” mutant SHP1-DA. (B) A volcano plot of the SHP1 PEPSI assay; the X axis represents the Log2 (difference) and the Y axis represents –Log10 (p-Value). The data was analyzed by Perseus_1.5.5.3 proteomics analysis tools. The significant hits identified in SHP1 PEPSI assay are labelled using different colors and shapes, and ranked based on the number of unique hit peptides. (C) THEMIS and SHP1-WT or DA mutant were electroporated into Jurkat cells, as indicated. Co-immunoprecipitation assays were performed to compare THEMIS binding affinity between SHP1-WT and SHP1-DA mutant. (D) A volcano plot of the PTP1B PEPSI assay; the X axis represents the Log2 (difference) and the Y axis represents –Log10 (p-Value). The data was analyzed by Perseus_1.5.5.3 proteomics analysis tools. The significant hits identified in PTP1B PEPSI assay are labelled using different colors and shapes, and ranked based on the number of unique hit peptides. (E) HEK293FT cells were transiently transfected with THEMIS-WT-FLAG, PTP1B-WT or PTP1B-DA, or SHP1-DA as control. Co-immunoprecipitation assays were performed to compare THEMIS binding affinity between PTP1B-WT, PTP1B-DA or SHP1-DA.

We applied the PEPSI platform to PTPN6/SHP1, a key PTP regulating T cell activation, to further identify its uncharacterized substrate(s) and characterize its function in TCR signaling. First, we generated a Jurkat T cell line with doxycycline-induced pupE expression. SHP1^WT^-PafA or SHP1^DA^-PafA fusion genes were then stably introduced into this cell line. Upon doxycycline stimulation and biotin addition, potential prey proteins which can associate with either WT or the substrate-trapping mutant form of SHP1 were biotinylated by PafA (Figure S1A). We enriched biotinylated protein, identified them by mass spectrometry, and plotted volcano map. The potential substrates demonstrated higher affinity with SHP1^DA^ substrate trapping mutant and were located in the right quadrant. We further ranked these candidate genes according to the number of unique peptides captured. THEMIS is on the top of the list (Figure 1B).

THEMIS (Thymocyte-expressed molecule involved in selection) is a T cell-specific protein which is critical for T cell development, especially CD4SP thymocytes maturation (Allen, 2009; Fu *et al*., 2009; Johnson *et al*., 2009; Kakugawa *et al*., 2009; Lesourne *et al*., 2009; Patrick *et al*., 2009). Furthermore, it is reported that the interplay between SHP1 and THEMIS is important to regulate the threshold of TCR signal transduction for positive and negative selection during T cell development (Choi *et al*., 2017a; Choi *et al*., 2017b; Fu *et al*., 2013; Gascoigne and Acuto, 2015; Gascoigne *et al*., 2016a; Gascoigne et al., 2016b; Mehta *et al*., 2018; Paster *et al*., 2015). To verify this interaction, immunoprecipitation experiments were performed in both Jurkat and HEK293T cell lines with co-expression of THEMIS and SHP1 WT or DA mutant, respectively. Results indicated that the substrate trapping mutant of SHP1 binds to THEMIS more tightly than WT (Figure 1C and Figure S1B-C), suggesting that THEMIS may be a novel substrate of SHP1.

To better assess the robustness of the PEPSI system and the specificity of the interaction between THEMIS and the SHP1 substrate trapping mutant, we further profiled the interactome of other tyrosine phosphatases such as SHP2 and PTP1B in the Jurkat T cell line. A previous study has revealed the association of THEMIS and SHP2 in SHP1 deficient mouse thymocytes (Mehta *et al*., 2018). In agreement with these observations (Mehta *et al*., 2018), ectopically expressed SHP2^DA^ or SHP1^DA^ bound to THEMIS with equivalent affinity, while PTP1B^DA^ did not (Figure 1D and Figure S1D). Biochemical immunoprecipitation experiments further confirmed no interaction between PTP1B^DA^ and THEMIS (Figure 1E and Figure S1E). Together, by harnessing the PEPSI platform, we demonstrated that THEMIS interacts more strongly with SHP1^DA^ substrate trapping mutant compared to SHP1 WT, implying an undisclosed substrate-enzyme regulatory mode between these two proteins.

### LCK phosphorylated THEMIS at Tyr34 site

Our PEPSI and follow-up biochemical verification suggest that the role of THEMIS in T cell development might rely on its tyrosine phosphorylation. Next, we addressed whether THEMIS undergoes tyrosine phosphorylation during T cell activation and the identity of the tyrosine kinase. We examined a series of tyrosine kinase(s), which are expressed in T cells, and found ectopically expressed LCK (Figure 2A) but not ZAP70 (Figure 2B), strongly phosphorylated THEMIS. To further pinpoint the specific site on which LCK acted, we individually mutated all 19 Y of THEMIS to F and compared the phosphorylation level change in the presence of LCK. Notably, the only site phosphorylated by LCK was the Tyr34 site (Figure 2C), which is highly conserved in multiple species possessing adaptive immunity (Figure 2D). Given the potential evolutionary significance of this sequence motif, we raised a polyclonal antibody to specifically detect the Tyr34 phosphorylation (pY34) of THEMIS. Whereas THEMIS-Y34F mutant could not be phosphorylated (Figure 2E), THEMIS 18YF mutant, in which only Y34 is intact, was readily recognized by the antibody (Figure 2F), indicating specificity of the antibody for pY34. Tyr34 resides in the CABIT1 domain of THEMIS. Domain truncation mutants of THEMIS that retains the CABIT1 domain could be phosphorylated by LCK, and this phosphorylation signal could be detected by the pY34-specific antibody (Figure S2A-B). In summary, these results demonstrated that Tyr34 of THEMIS is a bona fide substrate of non-receptor tyrosine kinase LCK.

**Figure 2.**
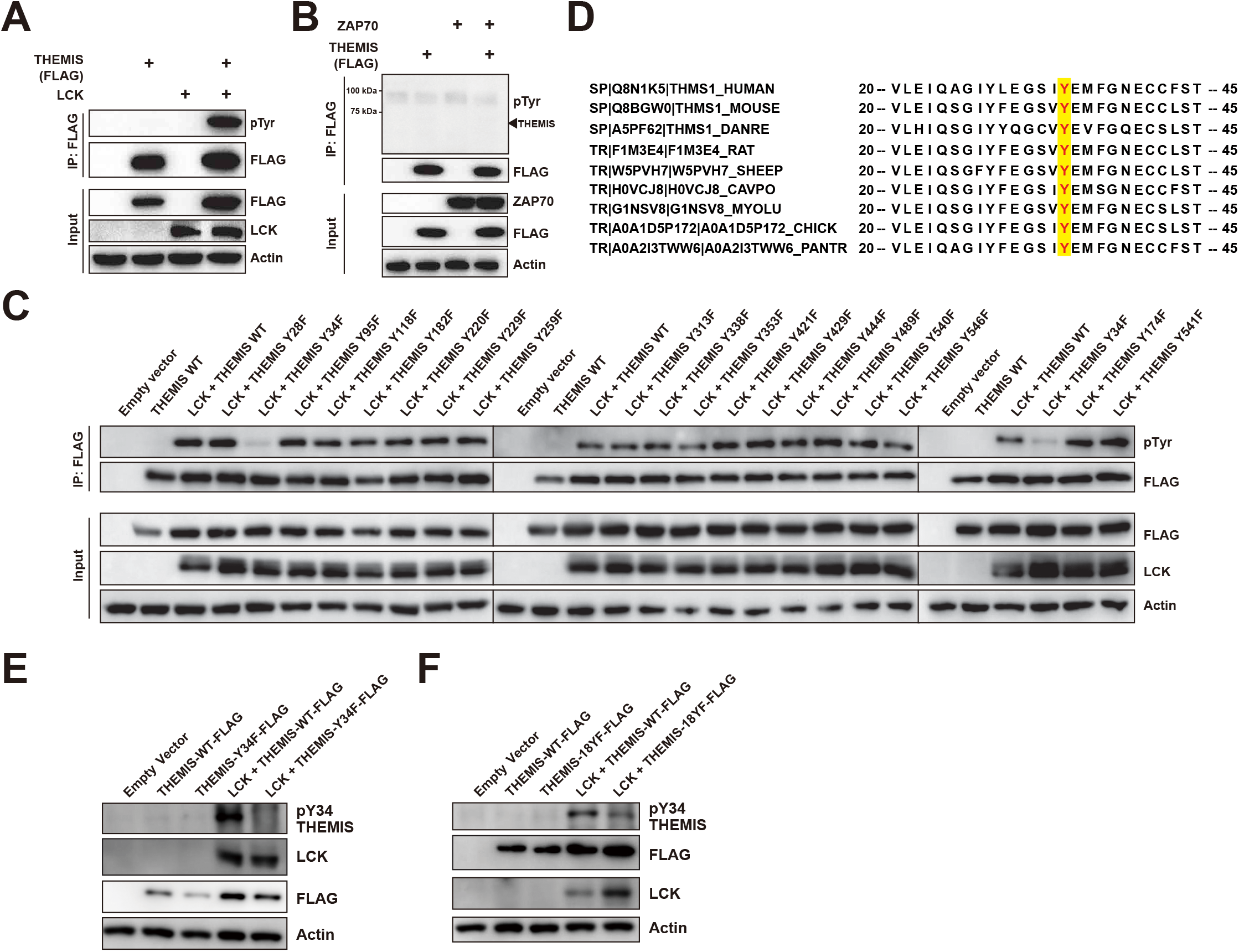
LCK phosphorylated THEMIS at Tyr34 site. (A-B) HEK293T cells were transiently transfected with THEMIS-WT-FLAG and LCK. (A) or ZAP70 (B) indicated LCK can phosphorylate THEMIS but not ZAP70. (C) HEK293T cells were transiently transfected with LCK and THEMIS-WT-FLAG or saturation phenylalanine mutation screen for all 19 of the tyrosine residues in the THEMIS protein. Western blotting indicated that LCK can phosphorylate THEMIS-WT but not the THEMIS-Y34F. (D) Sequence homology comparison of THEMIS Tyr34 site in different 9 species. (E-F) Experiments used extracts from HEK293T cells transiently transfected with LCK and THEMIS-WT, THEMIS-Y34F (a mutant bearing only the Y34F mutation), or THEMIS 18YF (a THEMIS mutant with its WT Y34 residue but with F substitutions at all 18 of its other Y residue positions). It indicated pY34 THEMIS antibody can specifically recognize the phosphorylation of THEMIS Tyr34 site.

### SHP1 dephosphorylated THEMIS at Tyr34 site after T cell activation

To further assess the notion that THEMIS is a novel substrate of tyrosine phosphatase SHP1, we reconstituted an *in vitro* phosphatase assay using phosphorylated pY34 peptide of THEMIS as substrate. Given the fact that intramolecular binding between N-terminal SH2 and PTP domains will induce an autoinhibited conformation that hinders phosphatase activity of SHP1 (Yang et al., 2003), we adopted E to A constitutively active form of SHP1 (Neel et al., 2003; O’Reilly et al., 2000; Wang et al., 2011) in our assay (Figure 3A). While constitutively inactive C to S form of SHP1 failed to dephosphorylate THEMIS at all, SHP1^EA^ protein robustly abrogated the phosphorylation of the pY34 peptide of THEMIS in a dose-dependent manner (Figure 3B). Furthermore, the Tyr34 phosphorylated THEMIS protein purified from LCK-expressing HEK293FT cells was dephosphorylated by SHP1^EA^, but not SHP1^CS^ protein *in vitro* (Figure 3C). In line with these results, Jurkat T cells stimulated with anti-TCR antibody illustrated overt tyrosine phosphorylation of THEMIS, peaking at 5 min post-stimulation (Figure 3D). Whereas genetic knockout (Figure 3D) or pharmacological inhibition (Figure S2C) of LCK totally abolished the phosphorylation signal, SHP1 deficiency in Jurkat cells further enhanced the intensity of pTyr34 phosphorylation on THEMIS (Figure 3D). Consistent with the previous mass spectrometry data, SHP1^DA^, the substrate trapping mutant form of the phosphatase robustly bound to WT THEMIS with enriched tyrosine phosphorylation, while its binding affinity was dramatically decreased in association with Y34F mutant form of THEMIS since it is no longer a substrate of SHP1 (Figure 3E). Taken together, results from these assays provided compelling evidence that kinase LCK and phosphatase SHP1 oppositely regulate the phosphorylation level of THEMIS at Tyr34 site during the T cell activation.

**Figure 3.**
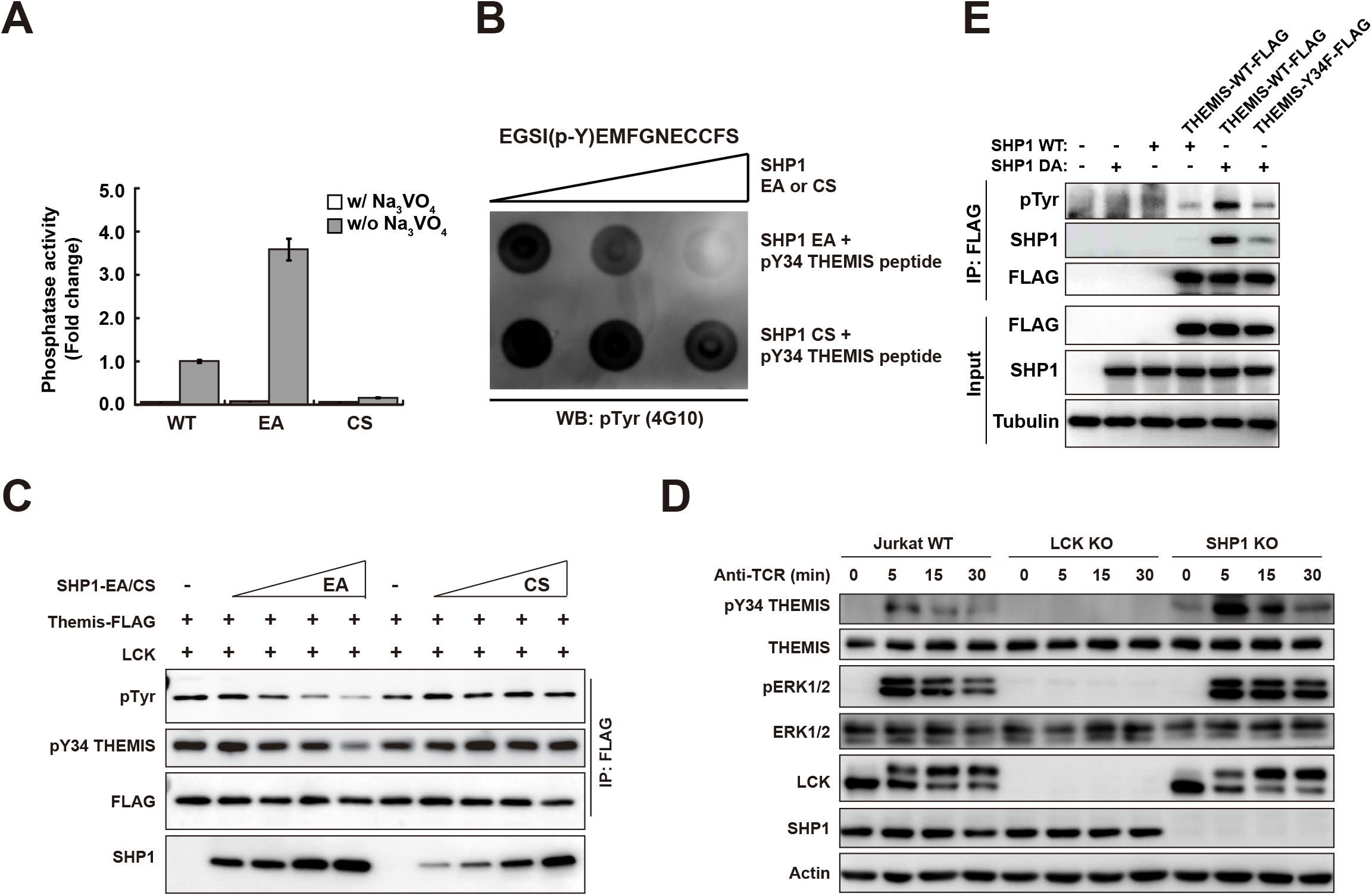
SHP1 dephosphorylated THEMIS at Tyr34 site after T cell activation. (A) *In vitro* DiFMUP assay which detect activity of protein phosphatase SHP1 showed that the constitutively activated SHP1-EA (E74A) mutant has the highest activity; the lowest activity was detected for the phosphatase-dead SHP1-CS (C453S) mutant. (B) Dot blot results indicating that the constitutively activated SHP1-EA mutant can dephosphorylate an artificial THEMIS p-Y34 peptide; the peptide sequence is shown above. (C) In vitro dephosphorylation assays which mixed the purified SHP1 protein from *E. coli* and the THEMIS protein affinity purified from HEK293T cells at 30 °C for 1 hour. Western blotting indicated that dephosphorylation of THEMIS Tyr34 is catalyzed by SHP1. (D) Immunoblotting analysis of the phosphorylation levels of THEMIS Tyr34 in Jurkat WT, LCK-KO Jurkat, and SHP1-KO Jurkat cells stimulated with an anti-TCR antibody (C305) for 0, 5, 15, and 30 min at 37 °C, demonstrating increased levels of phosphorylated THEMIS in SHP1-defective Jurkat cells. Additional immunoblotting against THEMIS, pERK1/2, ERK1/2, LCK, SHP1, and Actin (loading control). (E) HEK293T cells were transiently transfected with SHP1(WT or DA) and THEMIS (WT or Y34F). Western blotting suggest that THEMIS Y34F mutant weakens its binding to SHP1-DA comparing to THEMIS WT.

### CD4 T cell development, but not CD8 T cell development or peripheral expansion, depends on THEMIS Y34 phosphorylation

THEMIS is important in T cell maturation since the percentages of both CD4+ and CD8+ T cell development are substantially reduced at DP to SP stage in THEMIS knockout mice (Allen, 2009; Fu *et al*., 2009; Johnson *et al*., 2009; Kakugawa *et al*., 2009; Lesourne *et al*., 2009; Patrick *et al*., 2009). According to a compendium of single-cell transcriptomic data from multiple tissues in Mus musculus (Tabula Muris et al., 2018), *THEMIS* gene has a unique tissue expression pattern and is exclusively concentrated in thymus (Figure S3A). Among all populations of T cells, immature thymic T cells showed the highest expression level of THEMIS (Figure S3B).

Given the fact that the defect of SP T cell maturation in THEMIS^-/-^ mice could be fully rescued by additional T cell lineage-specific deletion of SHP1 (Choi *et al*., 2017b), as well as our previous data that highly supported an enzyme-substrate regulatory mode between SHP1 and THEMIS, we were very intrigued to further investigate the role of THEMIS phosphorylation at Tyr34 site in T cell development. To achieve this, we generated two C57BL/6J mice models by CRISPR technology (Hsu et al., 2014; Jinek et al., 2012): THEMIS knockout (THEMIS^-/-^) and Y to F knockin (THEMIS^Y34F/Y34F^) (Figure S3C). Immunoblotting analysis revealed normal protein expression of the phosphorylation-deficient mutant form of THEMIS in thymocytes (Figure 4A). Next, we performed flow cytometry analysis in THEMIS^+/+^, THEMIS^-/-^ and THEMIS^Y34F/Y34F^ mice to further assess the functional role of THEMIS phosphorylation (Tyr34) during T cell development. Consistent with the previous studies (Fu *et al*., 2009; Johnson *et al*., 2009; Kakugawa *et al*., 2009; Lesourne *et al*., 2009; Patrick *et al*., 2009), both sexes of THEMIS^-/-^ mice, compared to THEMIS^+/+^, exhibited an obvious decrease in number and percentage of CD4SP thymocytes (Figure 4B and Figure S3D), mature T (CD69^lo^TCRβ^hi^) thymocytes (Figure 4D and Figure S3F) and CD4 splenocytes (Figure 4E and Figure S3G), and to a lesser extent CD8SP thymocytes (Figure 4C and Figure S3E) and CD8 splenocytes (Figure 4F and Figure S3H). Similar to THEMIS^-/-^, phosphorylation-deficient THEMIS^Y34F/Y34F^ mice showed a slightly milder but significant decrease in number and percentage of CD4SP thymocytes (Figure 4B and Figure S3D), mature T (CD69^lo^TCRβ^hi^) thymocytes (Figure 4D and Figure S3F) and CD4 splenocytes (Figure 4E and Figure S3G), nevertheless have a normal maturation of CD8SP thymocytes (Figure 4C and Figure S3E) and CD8 splenocytes (Figure 4F and Figure S3H). Representative FACS histograms for three genotype mice are illustrated in Figures 4G-I. To further investigate the role of THEMIS phosphorylation (Y34) in CD8 T cell development, we crossed THEMIS^Y34F/Y34F^ and THEMIS^-/-^ mice with OTI TCR transgenic mice. Consistent with polyclonal TCR background, THEMIS Y34F mutation did not affect OTI-CD8 SP thymocytes development, whereas THEMIS deficiency substantially reduced OTI-CD8 SP thymocytes number (Figure 4J). Collectively, these *in vivo* results provide evidence to support the critical role of THEMIS phosphorylation at Tyr34 residue within the CABIT1 domain in T cell development and that abolishing this tyrosine phosphorylation regulatory potential results in a severe developmental alteration in CD4, but not CD8 T cell maturation.

**Figure 4.**
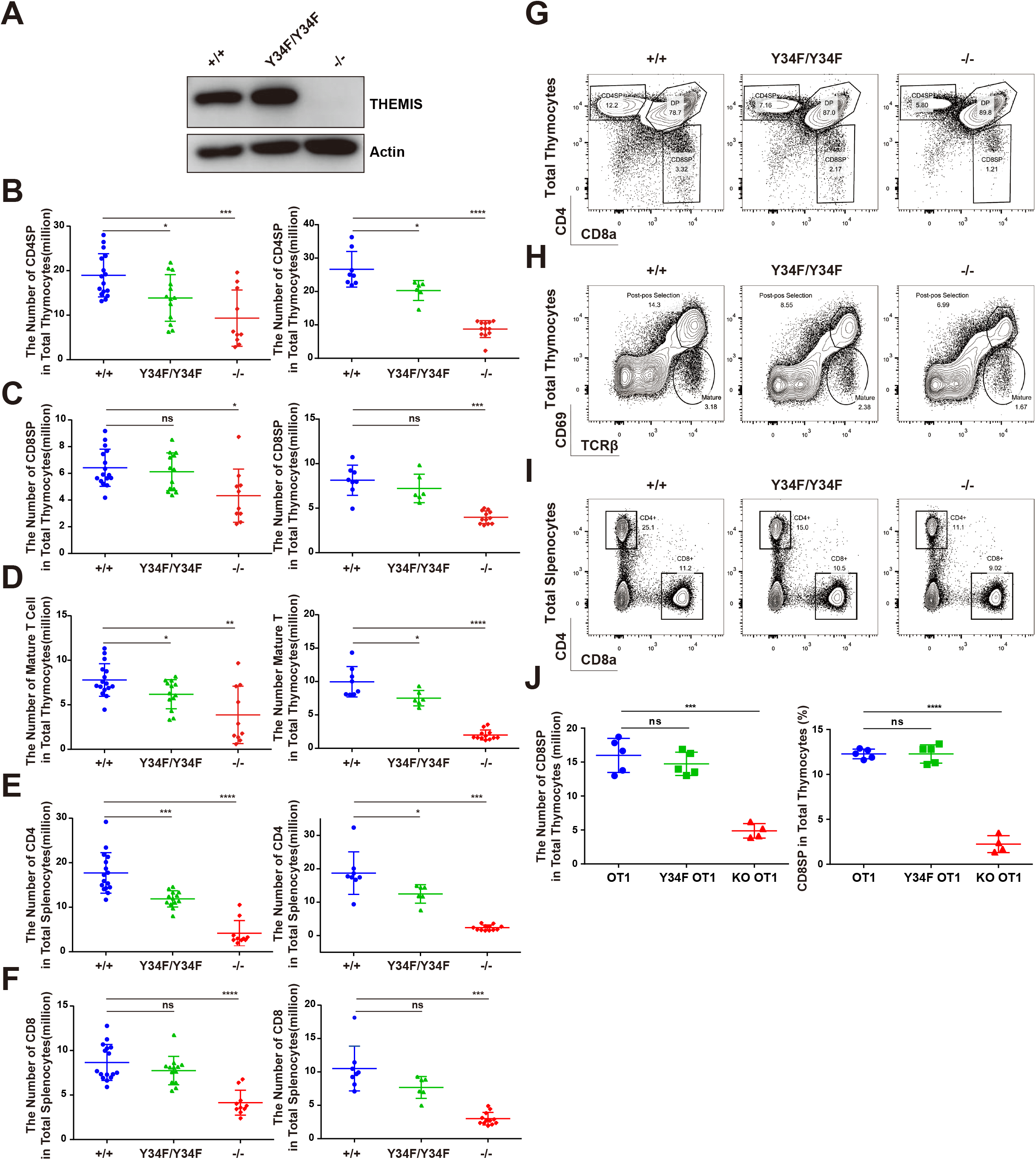
CD4 T cell development, but not CD8 T cell development or peripheral expansion, depends on THEMIS Y34 phosphorylation. (A) Western blot analysis of THEMIS and Actin (loading control) in total thymocytes isolated from THEMIS^+/+^, THEMIS^Y34F/Y34F^, and THEMIS^-/-^ mice. (B-F) Quantification analysis of CD4SP (B), CD8SP (C), and mature cells (D) in total thymocytes or CD4+ (E) and CD8+ (F) T cells among total splenocytes isolated from male mice (left) and female mice (right) of the three genotypes (male: n=16, 13, 10; female: n=8, 6, 12). * P<0.05, ** P<0.01, *** P<0.001, **** P<0.0001 (two tail unpaired t-test) (G-I) Representative flow cytometry histogram for analyzing number of CD4SP and CD8SP in both thymus and spleen tissues from THEMIS^+/+^, THEMIS^Y34F/Y34F^, and THEMIS^-/-^ mice. Mouse T cells (6∼7-week old) were stained with anti-CD4-Qdot605, anti-CD8-APC-Cy7, anti-TCRb-PerCP-Cy5.5, and anti-CD69-PE-Cy7 in thymocytes or CD4-PerCP-Cy5.5 and CD8-FITC in splenocytes for 1 hour on ice. (G) CD4 versus CD8 among total thymocytes, (H) CD69 versus TCRβ among total thymocytes, and (I) CD4 versus CD8 among total splenocytes. (J) Quantification analysis of CD8SP in total thymocytes (left, number; right, percentage) isolated from the OT1, THEMIS^Y34F/Y34F^ OT1 and THEMIS^-/-^ OT1 mice (n=5, 5, 4). * P<0.05, ** P<0.01, *** P<0.001, **** P<0.0001 (two tail unpaired t-test)

### Upon T cell activation, Tyr34 phosphorylated THEMIS played as a priming substrate to activate phosphatase activity of SHP1

The role of THEMIS in regulating the phosphatase activity of SHP1 is highly controversial. On the one hand, Gascoigne et al. demonstrated that THEMIS bound to SHP1 and enhanced phosphatase activity by increasing the phosphorylation level of SHP1 (Mehta *et al*., 2018). On the other hand, Love and his colleagues demonstrated that addition of THEMIS protein *in vitro* specifically inhibits phosphatase activity of SHP1, probably by stabilizing the oxidation of SHP1’s catalytic cysteine residue (Choi *et al*., 2017b). Considering Tyr34 of THEMIS is a *bona fide* substrate of SHP1 (Figure 3), we were very intrigued to compare the tyrosine phosphatase activity of SHP1 from mice expressing different forms of THEMIS. SHP1 protein was immunoprecipitated from mouse thymocytes, and its activity was evaluated with DiFMUP (6,8-difluoro-4-methylumbiliferyl phosphate) phosphatase assay. We chose DiFMUP as substrate for two reasons (Montalibet et al., 2005; Welte et al., 2005): 1) It has been well demonstrated to be an ideal substrate for fast, sensitive tyrosine phosphatase assays; 2) Compared to phospho-peptide substrate, DiFMUP is small, easily accesses the active site and has minimal impact on the conformation transition of the phosphatase. As shown in Figure 5A, TCR stimulated thymocytes from THEMIS^+/+^ mice exhibited significantly elevated phosphatase activity of SHP1. This enhancement was solely mediated by the presence of THEMIS, since a similar increase in activity was not observed in thymocytes from THEMIS^-/-^ mice (Figure 5A). Furthermore, phosphorylation of THEMIS at Tyr34 plays a key role in this phosphatase activity regulation, since a similar lack of increase in phosphatase activity was also observed in thymocytes from THEMIS^Y34F/Y34F^ phosphorylation-deficient mice (Figure 5A).

**Figure 5.**
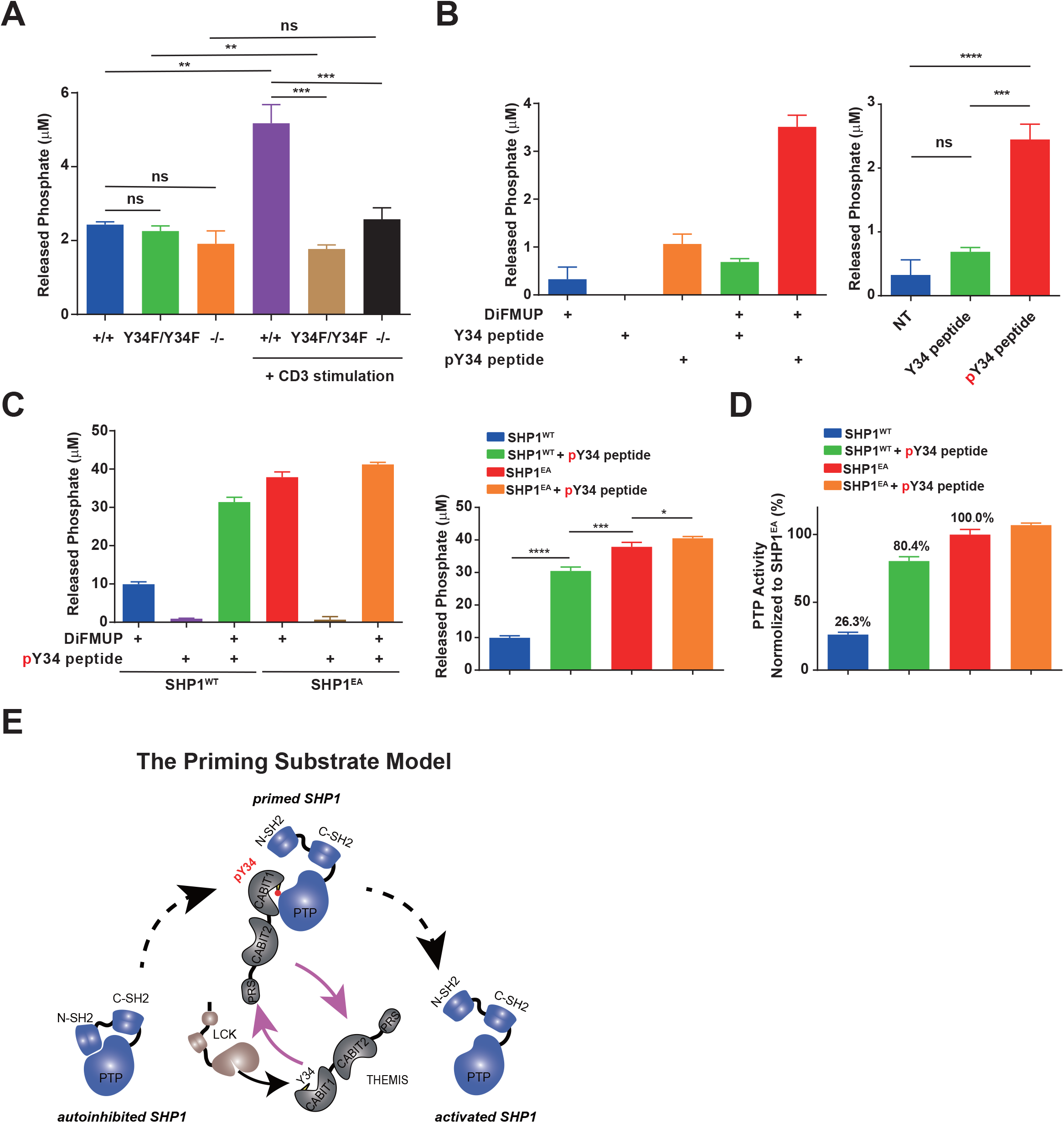
Upon T cell activation, Tyr34 phosphorylated THEMIS played as a priming substrate to activate phosphatase activity of SHP1. (A) Immnoprecipitated SHP1 from total thymocytes, unstimulated or stimulated with anti-mCD3 antibody for 2 mins, was incubated with DiFMUP at room temperature for 45 mins. The phosphatase activity was determined by malachite green phosphate assay. * P<0.05, ** P<0.01, *** P<0.001, **** P<0.0001 (two tail unpaired t-test) (B) SHP1 protein was immunoprecipitated from total thymocytes of THEMIS^-/-^ mice, and incubated with Y34 nonphosphorylated THEMIS peptide (EGSIYEMFGNECCFS) or Y34 phosphorylated THEMIS peptide (EGSIpYEMFGNECCFS). The phosphatase activity was determined by malachite green phosphate assay, with DiFMUP as substrate. * P<0.05, ** P<0.01, *** P<0.001, **** P<0.0001 (two tail unpaired t-test) (C-D) Recombinant SHP1^WT^ and SHP1^EA^ proteins were incubated with Y34 phosphorylated THEMIS peptide (EGSIpYEMFGNECCFS), followed by malachite green phosphate assay to determine phosphatase activity (C). The relative phosphatase activity was normalized to those of SHP1^EA^ proteins (D). * P<0.05, ** P<0.01, *** P<0.001, **** P<0.0001 (two tail unpaired t-test) (E) Tyr34 phosphorylated THEMIS acts as a priming substrate of SHP1 for its transition from autoinhibition to activation. During CD4 thymocyte development, Tyr34 of THEMIS is phosphorylated by LCK, and this phosphorylated motif serves as a docking site for recruitment and activation of SHP1.

All these results suggested that Themis Y34 phosphorylation can convert the activation status of SHP1 during T cell activation. To test this point, we synthesized both unphosphorylated and phosphorylated THEMIS peptides and tested their ability to regulate SHP1 activity. Notably, immunoprecipitated SHP1 from THEMIS^-/-^ mouse thymocytes showed prominently enhanced phosphatase activity triggered by Y34 phosphorylated THEMIS peptide, but not Y34 nonphosphorylated THEMIS peptide (Figure 5B), indicating the significance of Themis Y34 phosphorylation in modulating phosphatase activity of SHP1.

To further estimate the degree of SHP1 activity enhancement by pY34 THEMIS peptide, we performed the same experiment with recombinant SHP1 proteins, using SHP1^EA^ constitutive active mutant as positive control. As shown in Figures 5C and 5D, untreated SHP1^WT^ only possessed ∼26% phosphatase activity of SHP1^EA^, which is consistent with Figure 3A. However, addition of Y34 phosphorylated THEMIS peptide significantly enhanced the phosphatase activity of SHP1^WT^ by three folds, reaching ∼80% of SHP1^EA^. Interestingly, phosphatase activity of SHP1^EA^ was largely unchanged by Y34 phosphorylated THEMIS peptide (Figure 5D). These results highly suggested that Tyr34 of THEMIS, when phosphorylated, served as a priming substrate of SHP1 by exposing the PTPase site of the phosphatase and promoting the accessibility of the substrate.

### THEMIS also played phosphorylation-independent role in regulating CD8 T cell maturation and expansion

Interestingly, it is reported recently that THEMIS is required for the homeostasis of peripheral CD8 T cells and proliferative CD8 T cell responses to low-affinity pMHC aided by cytokines (Brzostek et al., 2020). A synergistic signal from both low-affinity pMHC and cytokine induces phosphorylation of Akt, metabolic changes and c-Myc transcriptional factor induction in CD8 T cells only in the presence of THEMIS (Brzostek *et al*., 2020). Given the previous data indicated that CD8SP thymocytes maturation was significantly dampened in THEMIS^-/-^, but not in THEMIS^Y34F/Y34F^ mice (Figure 4C and Figure S3E), we asked whether this Y to F mutation would affect peripheral CD8 T cell proliferative response upon stimulation. IL7 is required for the maintenance of peripheral T cell homeostasis (Fry et al., 2003; Sprent and Surh, 2011; Tamarit et al., 2013) and pro-inflammatory cytokine IL12 is indispensable for lymphopenia-induced proliferation (Goplen et al., 2016). Purified naïve CD8+ T cells from three genotypes were cultured in 50 ng/ml IL7 and IL12, stained with CFSE dye and analyzed by flow cytometry to monitor proliferative response. Consistent with the previous findings, THEMIS^-/-^ naïve CD8+ T cells showed a remarkable decrease in proliferation upon IL7 and IL12 stimulation compared to those from THEMIS^+/+^ mice (Figure 6A). Notably, no significant difference in cellular proliferation was observed between THEMIS^+/+^ and THEMIS^Y34F/Y34F^ mice (Figure 6A).

**Figure 6.**
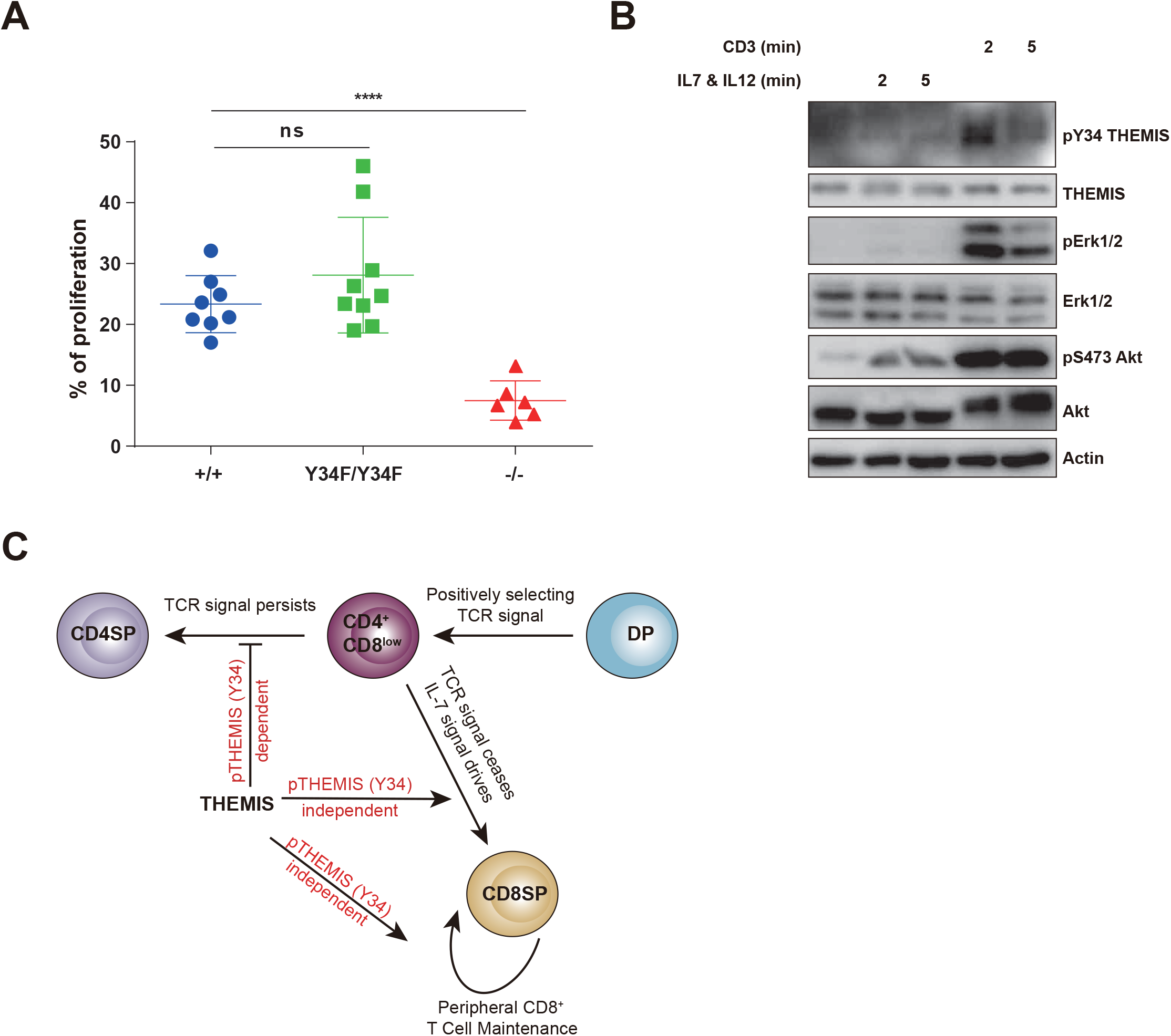
THEMIS also played phosphorylation-independent role in regulating CD8 T cell maturation and expansion. (A) Naïve CD8 T cell sorted by immunomagnetic negative selection system were labelled with CFSE dye and incubated with 50 ng/μl IL7 and IL12 for 7d. Cell proliferation were detected by CFSE staining analysis. Data were from n=8, n=9 and n=6 of three genotype mice. * P<0.05, ** P<0.01, *** P<0.001, **** P<0.0001 (two tail unpaired t-test) (B) Expanded mouse T lymphocytes were incubated with 50 ng/ul IL7 and IL12 or anti-CD3 antibody for 0 min, 2min and 5min. Western blot suggested IL7 and IL12 cytokine could not induce THEMIS Tyr34 phosphorylation while anti-CD3 stimulation can. Additional immunoblotting against THEMIS, pERK1/2, ERK1/2, pS473 Akt (positive control of successfully activation of IL7 and IL12) and Actin (loading control). (C) Working Model. Tyrosine phosphorylation-dependent manner: During CD4 thymocyte development, Tyr34 of THEMIS is phosphorylated by LCK, and this phosphorylated motif serves as a docking site for recruitment and activation of SHP1. Tyrosine phosphorylation-independent manner: During CD8 thymocyte development, IL-7 signaling dominates the generation of CD8 SP thymocytes as well as the homeostasis of peripheral CD8+ T cells without eliciting any phosphorylation on Tyr34 of THEMIS. Therefore, THEMIS functions in a phosphorylation-independent manner during these processes.

During T cell development, IL-7 signaling promotes CD8 thymocyte differentiation by silencing *Cd4* expression and by promoting reinitiation of *Cd8* gene transcription (Brugnera et al., 2000; Singer, 2002; Singer *et al*., 2008). Retrospect the experiment which explores the function of THEMIS in T cell development, we demonstrated that Y34F mutation of THEMIS, unlike THEMIS KO, has no significant impact on CD8 T cell maturation (Figures 4C and 4F). Concurrently, the experiment evaluating peripheral CD8 T cell proliferation after IL-7 & IL-12 co-stimulation also illustrated that THEMIS itself, but not the phosphorylation state of Tyr34 of THEMIS, is involved in peripheral CD8 T cell expansion and homeostasis (Figure 6A). Therefore, we speculated that THEMIS might exert phosphorylation-independent role in IL-7-mediated CD8 T cell maturation and peripheral proliferation. To investigate this hypothesis, we compared the phosphorylation status of THEMIS at Tyr34 in mouse lymphocytes by stimulation with either anti-CD3 or IL7 & IL12 co-administration. As expected, anti-CD3 treatment elicited robust elevation in protein phosphorylation on THEMIS, Erk1/2 and AKT (Figure 6B). Whereas AKT phosphorylation was enhanced upon IL7 & IL12 co-stimulation, we did not observe phosphorylation of THEMIS at Tyr34 (Figure 6B), supporting the notion that tyrosine phosphorylation regulation of THEMIS is not required for CD8 T cell peripheral proliferation.

## Discussion

The capability to define substrate specificity of protein tyrosine phosphatases is critical to gain an understanding of the distinct function of these enzymes. Harnessing insights from crystal structure and mutational analysis, Tonks and his colleagues established a mutant form of PTP that maintains its high affinity to substrate(s) but does not catalyze dephosphorylation reaction effectively, thereby converting the active enzyme into a “substrate trap” (Flint *et al*., 1997). Mutation of the invariant catalytic acid (Asp) yields such a substrate-trapping mutant. The fact that this Asp residue is conserved across almost all members of the PTP family implies that this mutational manipulation may offer a uniform strategy to identify physiological substrates of PTPs. However, this technology has its own limitations: 1) low-throughput; 2) the enhanced binding affinity between enzyme and substrate is not always strong enough for substrate identification through conventional biochemical approach (e.g. immunoprecipitation). To overcome these limitations for substrate identification in the PTP field, we took advantage of a newly invented proximity labeling technology called PUP-IT (Liu *et al*., 2018), which is ideal for weak binding detection, and integrated it with substrate trapping strategy to develop a systematic proximity labeling technology named PEPSI. This improved strategy will almost certainly facilitate comprehensive exploration of the over a hundred protein tyrosine phosphatases in the human genome and should help identify their potentially thousands of substrates deployed ubiquitously throughout cellular signaling and metabolism.

Illustrating the utility of the PEPSI platform, we focused our efforts on a protein tyrosine phosphatase known to function in both innate and adaptive immunity - SHP1 (Abram and Lowell, 2017). Interestingly, PEPSI revealed that the top candidate protein of SHP1’s dephosphorylation activity is THEMIS, which has recently gained major attention regarding its critical role in T cell development. By performing a series of biochemical analyses, we provided compelling evidence to confirm that LCK and SHP1 catalyzed opposing phosphorylation and dephosphorylation of THEMIS at Tyr34 respectively (Figures 2 and 3). Although several groups have reported the physical interaction between THEMIS and SHP1, mainly through their CABIT1 and PTP domains respectively(Choi *et al*., 2017b), we are the first to demonstrate the phosphatase-substrate action mode to reveal the details of the inter-regulation of these two proteins.

Whether THEMIS regulates SHP1’s activity positively or negatively is a key, but unsolved, question in THEMIS research. Currently, there are two models regarding the role of THEMIS in T cell selection (Choi *et al*., 2017a). Gascoigne and his colleagues illustrated that THEMIS enhances SHP1 phosphatase activity by increasing its phosphorylation, therefore proposing a negative regulatory role of THEMIS in TCR signal strength during thymic selection (Fu *et al*., 2013; Mehta *et al*., 2018). However, Love and his colleagues reported that THEMIS is a negative regulator of SHP1 phosphatase activity through reactive oxygen species (ROS)-dependent regulation of SHP1, therefore playing a positive role in TCR signaling and positive selection (Choi *et al*., 2017b). Both groups adopted the same malachite green phosphatase assay to measure SHP1’s activity, with the difference that the former group used enriched proteins from ex vivo thymocytes and the latter group used purified proteins. Both groups agree that THEMIS is essential in moderating the phosphatase activity of SHP1 but reach contradictory conclusions regarding either a positive or negative mode of regulation. Our data supports THEMIS as a positive regulator of SHP1’s activity and therefore plays a negative role in TCR signaling and positive selection. By performing a standard ex vivo tyrosine phosphatase DiFMUP assay to measure the PTP activity associated with immunoprecipitated proteins directly, we found that SHP1’s activity was significantly reduced in activated thymocytes from both THEMIS^-/-^ and THEMIS^Y34F/Y34F^ mice, relative to WT mice (Figures 5A-D). Most importantly, we demonstrated that THEMIS is a bona fide substrate for SHP1. Therefore, excessive purified THEMIS, which was added into the in vitro dephosphorylation assay, may compete with the phospho-peptide substrate provided by malachite green phosphatase assay for binding to the phosphatase, leading to the decrease of the “apparent phosphatase activity”, observed in Love’s studies (Choi *et al*., 2017b).

In addition, our results shed more insight into the mechanism of how THEMIS exerts a positive function is regulating SHP1’s activity. THEMIS, mainly through its CABIT1 domain, has been reported to bind to the phosphatase domain of SHP1, although molecular details remain lacking (Choi *et al*., 2017b). By performing both dephosphorylation and substrate trapping assays, we demonstrated that phosphorylated Tyr34 of THEMIS, which resides in the middle of the CABIT1 domain, is important for association with SHP1, prior to its dephosphorylation. Given that SHP1’s activity is severely decreased in THEMIS-deficient thymocytes, tyrosine phosphorylated THEMIS may serve as a “priming substrate” to recruit SHP1 and open its active site thus enhancing phosphatase activity (Figure 5E and Figure S4). In the absence of THEMIS, SHP1 likely stays in a closed conformation with its phosphatase domain occluded by its own N-SH2 domain, thus rendering the enzyme inactive (Abram and Lowell, 2017; Yang *et al*., 2003). Primed SHP1 dephosphorylates THEMIS, discharges THEMIS from the active site and is ready for other potential substrates in downstream TCR signaling. Notably, this model also explains much easier access of ROS to the active site of SHP1, resulting in cysteine oxidation and phosphatase inactivation in the presence of THEMIS.

Based on experimental observations obtained so far, the highly complicated process of T cell development could be best explained by the kinetic signaling model, proposed by Alfred Singer (Singer *et al*., 2008). According to this model, TCR signaling is continuously required for CD4 cell maturation, from DP to SP. Upon TCR stimulation, THEMIS is phosphorylated at Tyr34 and this tyrosine phosphorylation is crucial for the conformational transition of SHP1 from inactive to active. Activated SHP1 could thus fine-tune the TCR signaling for CD4 maturation in a negative manner. Consistently, compared to WT, THEMIS Tyr34 phosphorylation-deficient mice show profoundly decreased phosphatase activity (Figure 5A), which likely increase negative selection of CD4 SP thymocytes (Figure 4 and Figure S3). During CD8 maturation, however, TCR signaling is switched off in the later stage and replaced by cytokine signal (Singer *et al*., 2008). That IL7R blockade leads to abrogation of CD8 T cell development emphasizes the essentiality of IL7 signaling in CD8 T cell maturation (Yu et al., 2003). Interestingly, IL7 was completely unable to elicit tyrosine phosphorylation of THEMIS (Figure 6B). In line with this result, THEMIS Tyr34 phosphorylation-deficient mice, unlike THEMIS^-/-^ mice, show no defects in both CD8 SP T cell maturation and peripheral CD8 T cell expansion (Figure 4 and Figure S3).

We propose the following working model (Figure 6C): 1) THEMIS is required for both CD4 and CD8 T cell maturation. 2) THEMIS exerts its function in both tyrosine phosphorylation-dependent and -independent manner, depending on the stimuli. During CD4 thymocyte development, Tyr34 of THEMIS is phosphorylated by LCK, and this phosphorylated motif acts as a *“Mamba’s Fang”* to bite the PTP domain of SHP1 and convert its phosphatase activity from basal level to nearly fully activated level (Figure 5E and Figure S4). Primed SHP1 inhibits downstream TCR signaling and ensures an appropriate negative selection (Figure 5E). Deleting SHP1 or blocking THEMIS Y34 phosphorylation results in failure to prime SHP1’s activity, leading into enhanced negative selection (Fu *et al*., 2013). During CD8 thymocyte maturation, IL-7 signaling has a dominate role in the generation of CD8 SP thymocytes, and in this context THEMIS functions as in a phosphorylation-independent manner (Figure 6C), which also plays an important role in homeostasis expansion of periphery CD8 T cells. 3) A delicate feedback loop among TCR-LCK-pTyr34 THEMIS-SHP1 to ensure proper signaling duration and intensity is required for T cell selection. We strongly believe that the basic insights of this study will uncover the regulatory mechanism of THEMIS and SHP1 during T cell maturation and function, and its promising clinical purpose and potential implications will further optimize the engineered T-cell immunotherapy (Sun et al., 2020).

## Methods

### Cell culture

HEK293T cell line was cultured in Dulbecco’s modified Eagle’s medium (DMEM, Cellgro, 10-013-CV) containing with 10% FBS (PAN, ST30-3302), 1% penicillin and streptomycin (100×, TransGen, FG101-01). Jurkat cell line was cultured in RPMI 1640 medium (Cellgro, 10-040-CVR) supplemented with 5% FBS, 1% penicillin and streptomycin.

Primary mouse T cells (stimulation with anti-mCD3 (BioXcell, BE0001-1) and anti-mCD28 (BioXcell, BE0015-1) antibodies coated on plate) were cultured in supplementary DMEM medium (sDMEM) containing 10% FBS, 1% penicillin and streptomycin, 1× MEM NEAA (100×, Gibco, 11140050), 50 µM mercaptoethanol (Gibco, 21985023), 1 mM sodium pyruvate (Gibco, 11360070), 1× GlutaMAX™ (100×, Gibco, 35050061) and 25 mM HEPES (Gibco, 25-0691-82). After anti-CD3 and anti-CD28 antibodies stimulation 36-48h, cells were transferred into new plate and cultured in supplementary DMEM medium containing human IL2 cytokine (Novoprotein, GMP-CD66). Cells were cultured at 37°C in an atmosphere of 5% CO_2_.

### Plasmids

Mammalian expression plasmids used in this study were: pJ3-SHP1-WT (Addgene, 8572), pJ3-SHP1-D419A, pMT2-PTP1B-WT/D181A, pcDNA3.1-SHP2-D425A, pEF6-LCK and pEF6-ZAP70. PUP-IT related plasmids: pHR-SHP1-WT-PafA-Myc-IRES-EGFP and pHR-SHP1-D419A-PafA-Myc-IRES-EGFP, pHR-PTP1B-WT-PafA-Myc-IRES-EGFP and pHR-PTP1B-D181A-PafA-Myc-IRES-EGFP, pHR-SHP2-D425A-PafA-Myc-IRES-EGFP. pcDNA6-THEMIS-Flag was kindly provided by Prof. Guo Fu, Xiamen University, P.R. China. Using this plasmid as template, we constructed 19 single YF mutants and 18YF mutant (only retain Tyr34) of THEMIS (primer sequences in Table 1) by mutagenesis (QuikChangeTM Site-directed Mutagenesis Kit, 200518). We also subcloned THEMIS truncation mutants 1-260, 260-493, 1-493, 493-641 and 260-641 by homologous recombination (Beyotime, D7010S) into pEF6-Myc vector (primer sequences in Table 2).

**Table 1:**
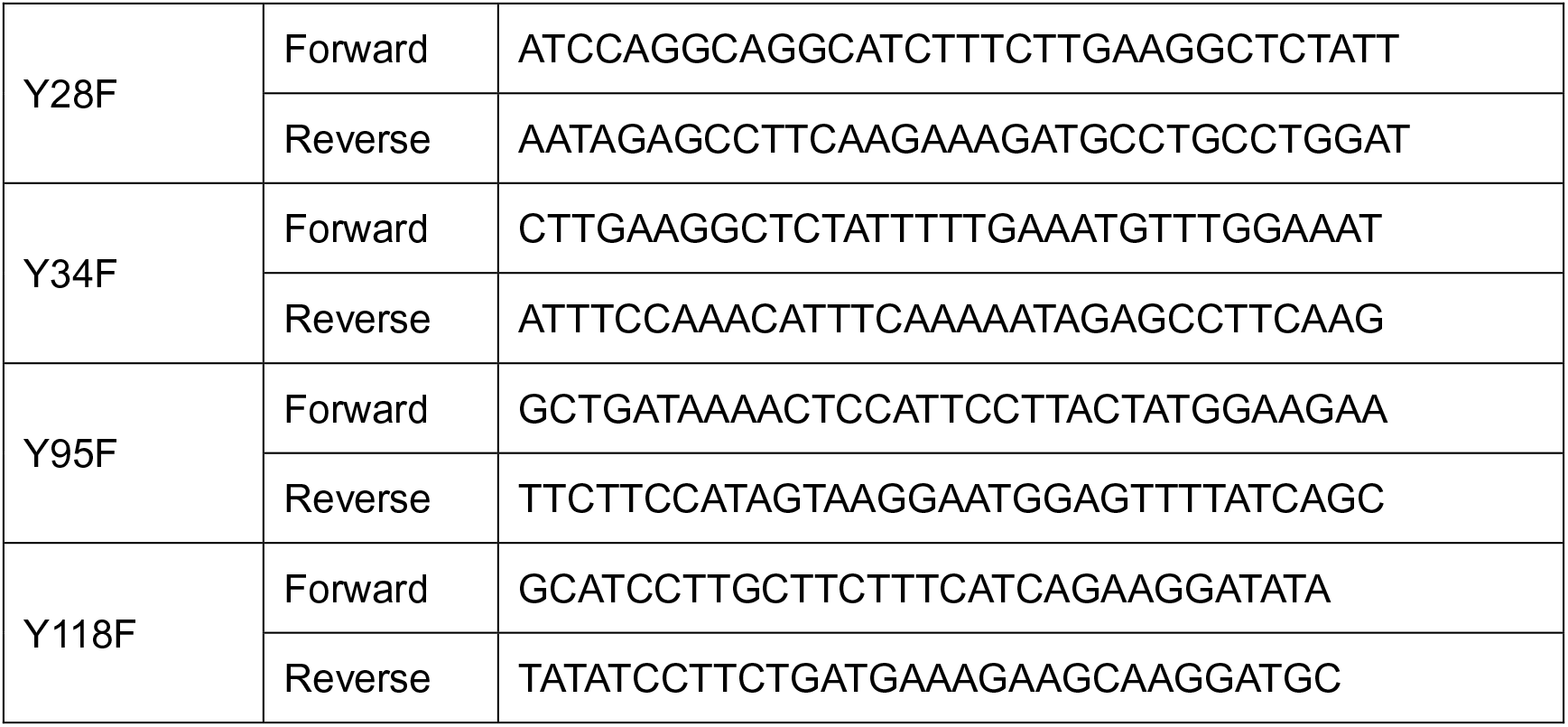
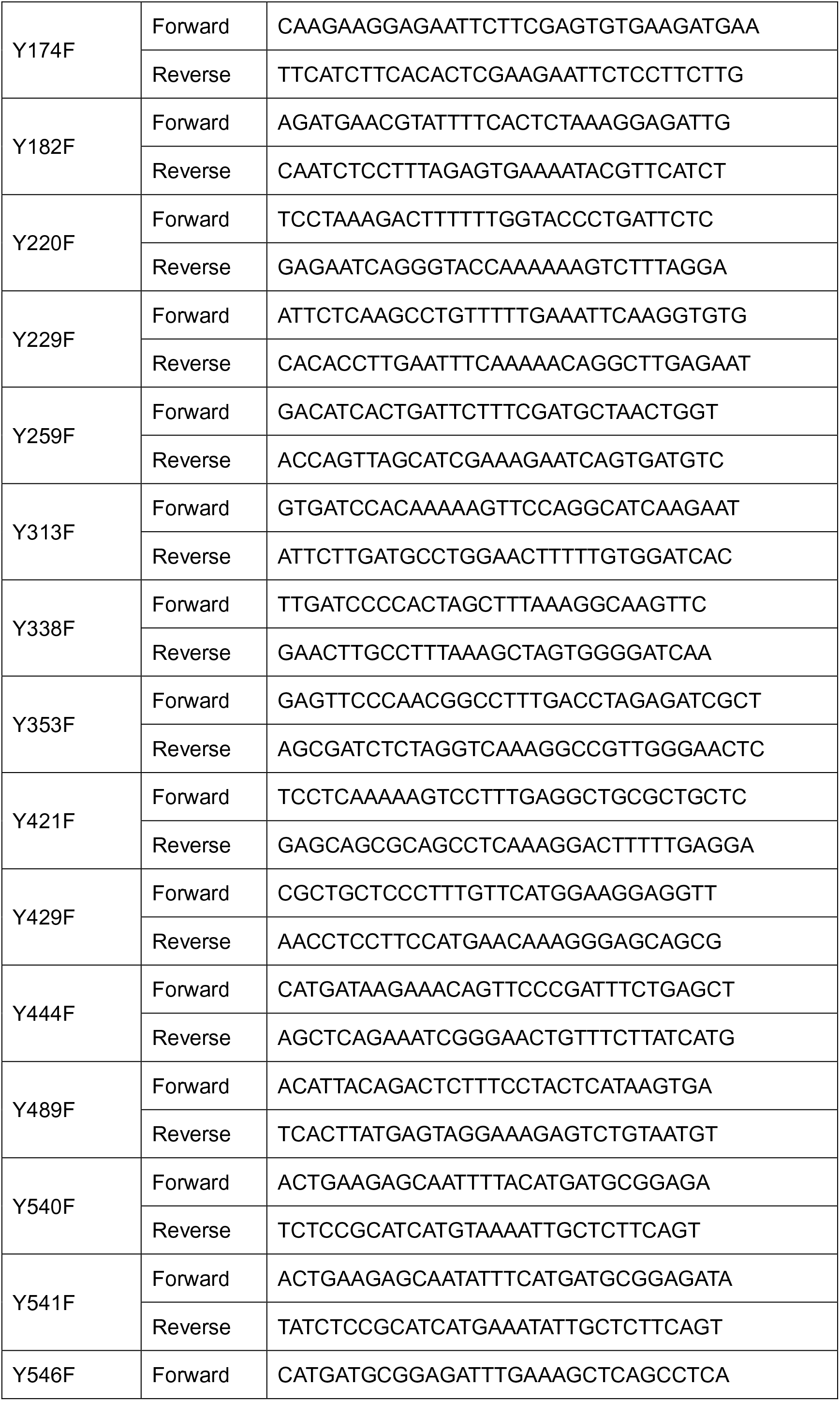
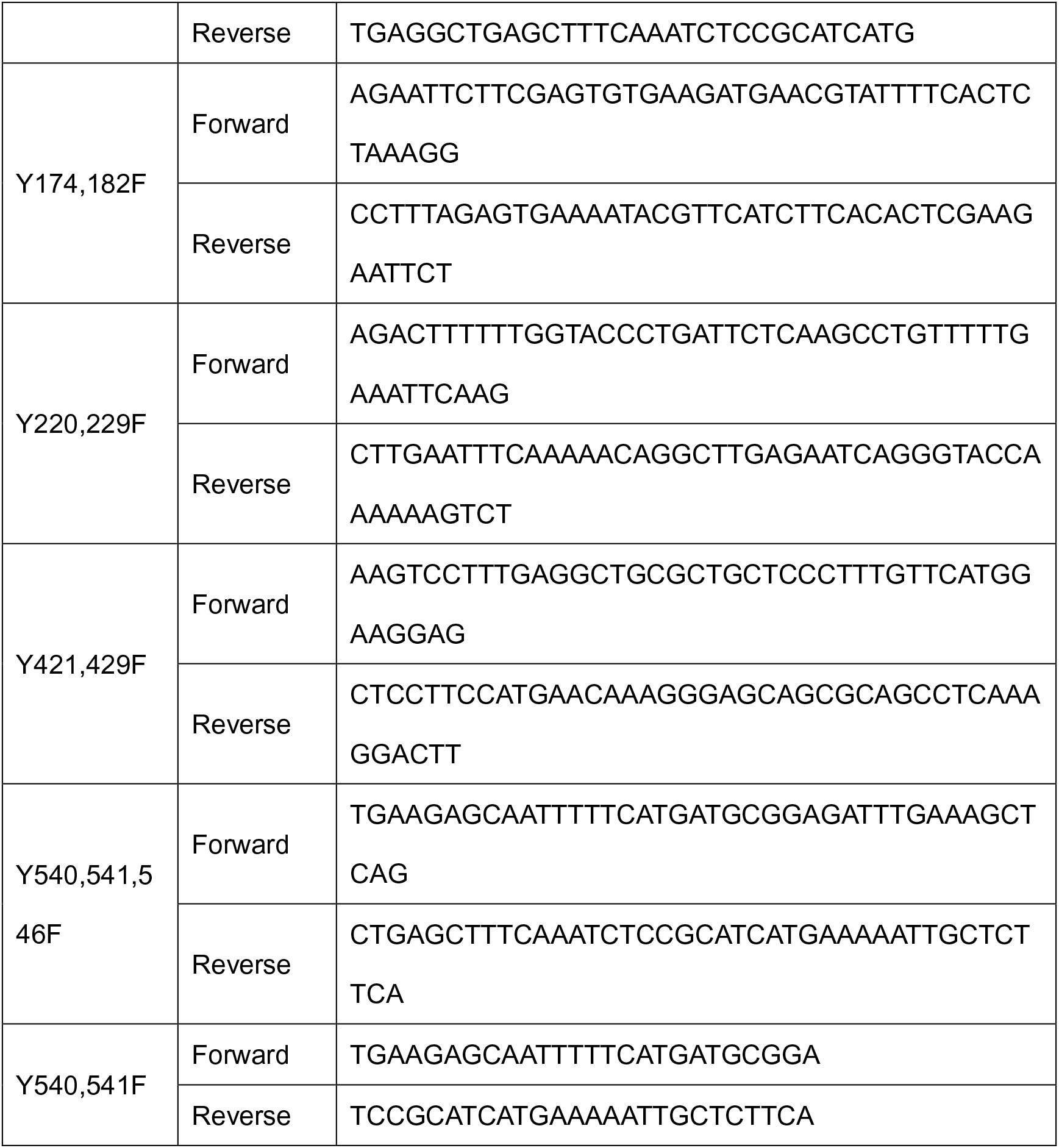
THEMIS Y to F mutants

**Table 2:**
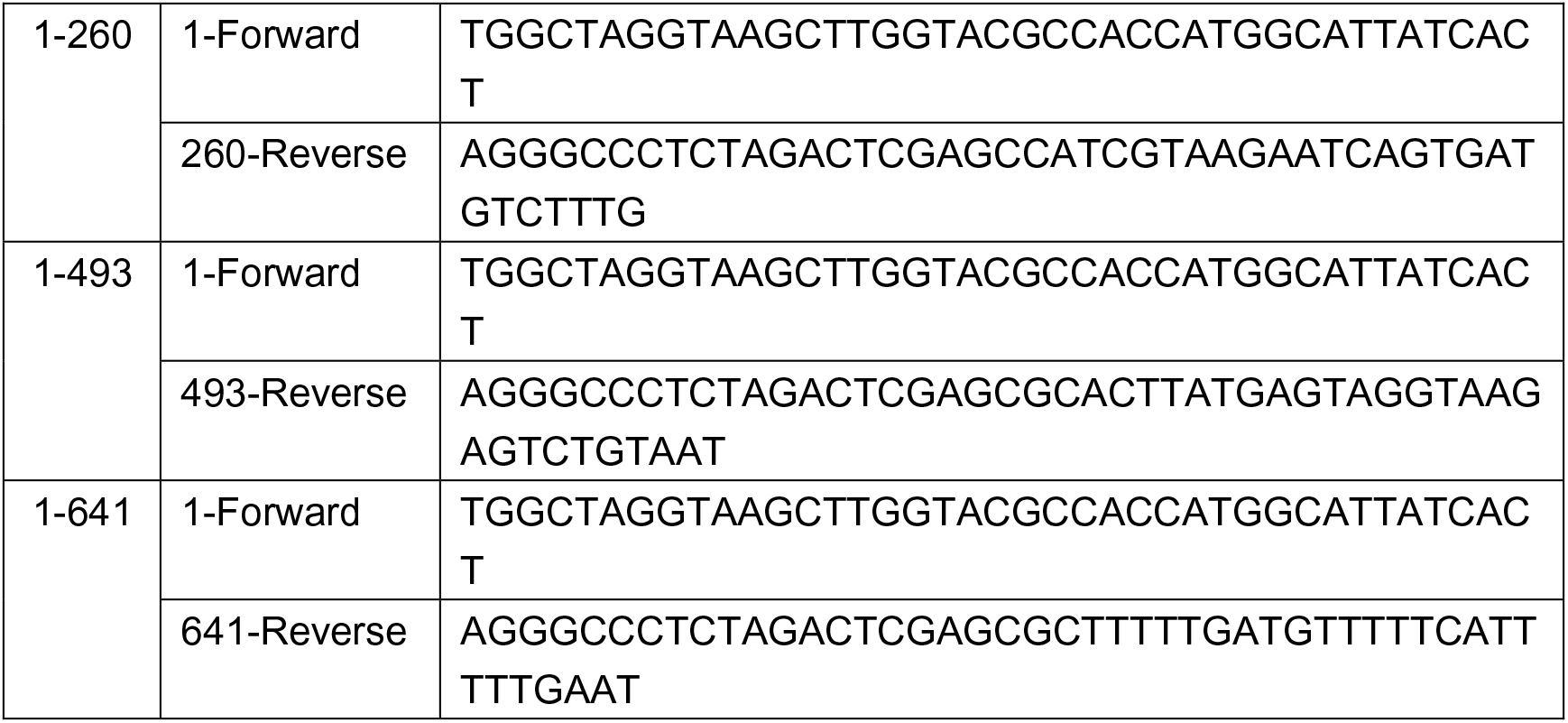
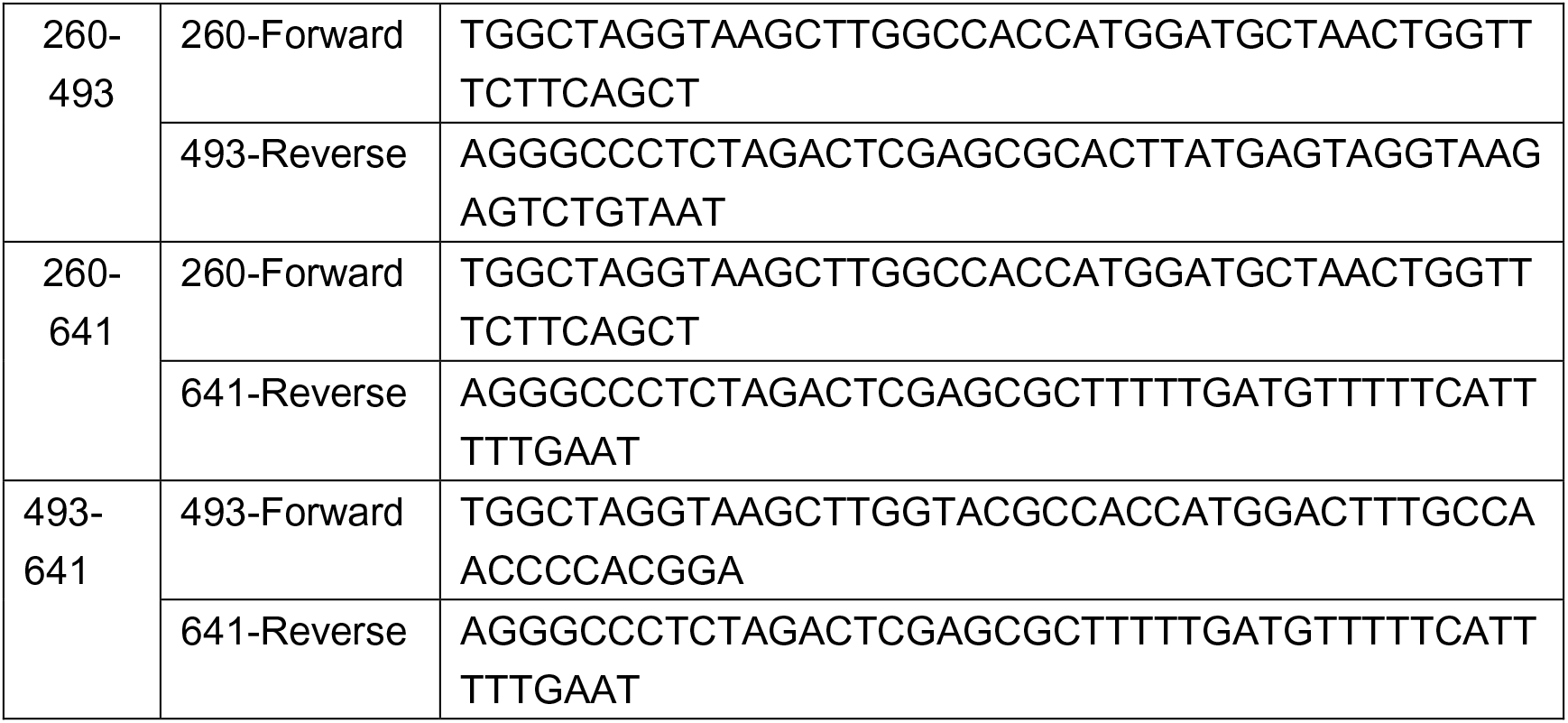
Themis truncated mutants in PEF6 vector

### Cell transfection and infection

Transient transfection was performed with Mirus (TransIT-2020, Mirus Bio) in 6-well plate. When cells were >80% confluence, we prepared TransIT-2020 reagent and DNA mixture with medium: 2.5 µg plasmid DNA and 5 µl TransIT-2020 reagent were added into 250 µl of Opti-MEMI Reduced-Serum Medium (Gibco). After adding the mixture into each well, cells were cultured about 24 hours and then harvested and lysed for following immunoprecipitation or immunoblotting assays.

Stable cell lines were established by lentiviral infection. Lentivirus was produced in HEK293T cell by transfecting plasmids (gene containing plasmids, virus packaging plasmid pCMVdR8.91 and pMD2.G) at a ratio of 10:10:1. After 48-72 hours, supernatants were harvested and passed through 0.45 μm filters to remove cells. Virus were then added into host cells about 48-72 hours. Infected cells were sorted by fluorescence (BD Aria) or selection drug to generate stable cell line. The effectiveness of infection was confirmed by flow cytometry or immunoblotting analysis with according antibody.

### CRISPR-Cas9 system for gene knockout

We applied CRISPR-Cas9 system to generate SHP1-knockout Jurkat cell lines. sgRNAs were designed from the website (http://www.e-crisp.org/E-CRISP/) and were subcloned into PX330-Cas9-GFP plasmid (Table 3). The recombinant plasmids were transfected into Jurkat cell line by electroporation. After 24-48 h, we sorted single clone GFP positive cells into 96-well U-bottom plate by flow cytometry (BD Aria). It takes ∼3 weeks for cell re-population, followed by immunoblotting analysis validation. LCK-knockout Jurkat cell line was kindly provided by Prof. Arthur Weiss (UCSF)

**Table 3:**
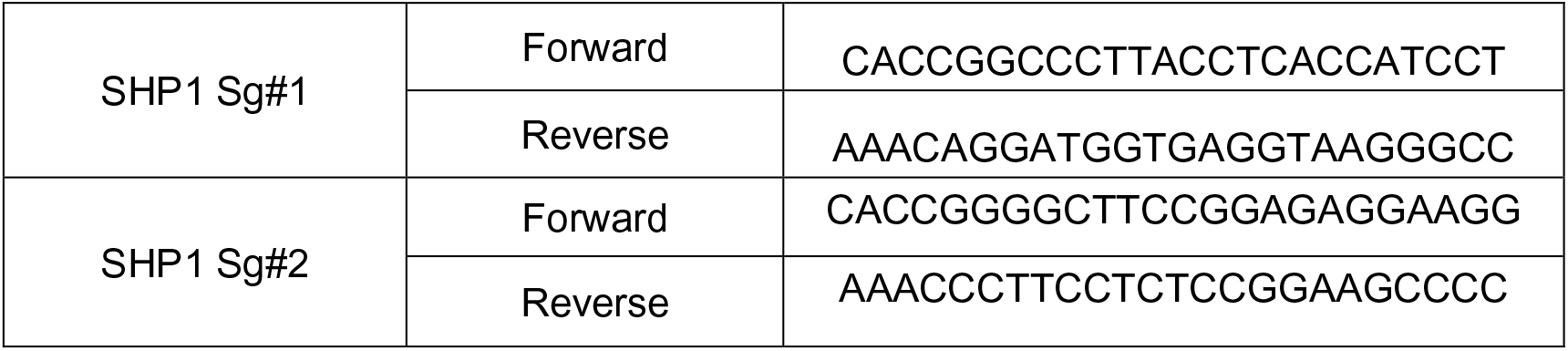
SHP1 sgRNA cloned in PX330

### Immunoblotting and immunoprecipitation

Cell lysates were extracted by 1% NP40 lysis buffer (20 mM Hepes pH7.5, 150 mM NaCl, 1% NP40, 1 mM Na_3_VO_4_) at 4°C for 30 minutes. Total protein concentration was determined by Bradford (Transgen, GP301-01) or BCA (Beyotime, P0010) assay.

For immunoblotting, cellular proteins were harvested, separated by SDS-PAGE and transferred onto nitrocellulose membranes. Membranes were blocked in 2.5% BSA in TBST (TBS/Tween 20: 20 mM Tris HCl, pH 7.5, 50 mM NaCl, and 0.1% Tween 20) for 1 hour at room temperature on a shaker and incubated with primary antibody at 4°C overnight. Proteins were detected with horseradish peroxidase (HRP)-conjugated secondary antibodies (Jackson Laboratory) and ECL (Pierce).

For immunoprecipitation, precleared cell extracts were incubated with the indicated antibody for 4 hours at 4°C with rotation followed by 1 hour of pull-down by 1:1 protein A/G agarose beads (GE, 17061801, 17127901) or FLAG/Myc beads (FLAG resin: Sigma, A2220; Myc resin: Sigma, E6654). Immunoprecipitates were washed with lysis buffer three times before electrophoresis.

### Protein purification

SHP1-WT, SHP1-E74A, SHP1-D419A and SHP1-C453S were subcloned into pET28a plasmid and were transformed into BL21 competent *E.coli* for protein expression and purification. Final concentration of 0.2 mM IPTG was used for induction at 16 °C for 20 hours. Harvested cells were resuspended in lysis buffer (50 mM Tris 8.0, 200 mM NaCl), and lysed with a pressure cell press. Supernatants were obtained by centrifugation and mixed with Ni-NTA resin to purify proteins on a gravity column. Finally, proteins were washed by washing buffer (50 mM Tris 8.0, 200 mM NaCl, 20 mM imidazole) and eluted by elution buffer (50 mM Tris 8.0, 200 mM NaCl, 100 mM imidazole). The purified proteins were added with glycerol to a final concentration of 10% and then were stored at −80 °C.

### *In vitro* DiFMUP phosphatase assay

*In vitro* DiFMUP assay was performed to evaluate the phosphatase activity of recombinant proteins SHP1-WT, SHP1-E74A and SHP1-C453S. Each reaction was reconstituted with 10 μl protein (2 ng/μl), 10 μl 100 μM DiFMUP (Thermo, D6567) and 80 μl assay buffer (50 mM Hepes PH 7.0, 150 mM NaCl, and freshly added 0.1 % BSA and 2 mM DTT) in white bottom 96-well plate at room temperature for 30 mins. The results were documented by microplate reader at excitation/emission of 358/450 nm.

### *Ex vivo* Malachite Green phosphate assay

SHP1 proteins were immunoprecipitated from thymocytes (50 million cells per sample) of THEMIS^+/+^, THEMIS^Y34F/Y34F^ and THEMIS^-/-^ mice in lysis buffer containing 5 mM DTT(Mehta *et al*., 2018), followed by 1x wash in Tris-NaCl buffer and 2x washes in the phosphatase buffer (25 mM HEPES [pH 6.5], 150 mM NaCl, 0.1 mM EDTA, 0.01 % Brij 35). Immunoprecipitates were resuspended in 150 µl phosphatase buffer, with addition of 2 µl DiFMUP (5 mM) to allow dephosphorylation reaction at room temperature for 45 mins. 40 µl supernatant for each reaction was then diluted with the phosphatase buffer at 1:1 ratio in 96-well plate, with addition of 20 µl malachite green dye mix (Sigma, MAK307-1KT) for released phosphate detection. Molar concentration of released phosphate in each sample was calculated by phosphate standards provided with the kit.

For *in vitro* substrate priming assay, immunoprecipitated SHP1 proteins from thymocytes (200 million cells) of THEMIS^-/-^ mice were divided into five equal portions and cultured with 2 µl (2 mM) THEMIS peptide (EGSIYEMFGNECCFS) or pY34 THEMIS peptide (EGSIpYEMFGNECCFS), as indicated, for 60 mins at room temperature. Then samples were added 2 µl DiFMUP (5 mM) to allow dephosphorylation reaction at room temperature for 60 mins. The released phosphate was detected by Malachite green assay.

For *in vitro* recombinant SHP1 phosphatase assay, 150 ng SHP1^WT^ and SHP1^EA^ were first cultured with 1 µl (2 mM) pY34 THEMIS peptide (EGSIpYEMFGNECCFS) for 15 mins at room temperature. Then samples were added 2 µl DiFMUP (5 mM) to allow dephosphorylation reaction at room temperature for 10 mins. The released phosphate was detected by Malachite green assay.

### Dot blot assay

300 ng pY34 THEMIS peptide (EGSIpYEMFGNECCFS) was incubated with 0.5, 1, 2 μg purified SHP1^EA^ or SHP1^CS^ protein at 30 °C for 1 hour for dephosphorylation. Sample were added SDS (final concentration 2%) and then heated at 95 °C for 5 mins to stop the reaction. 5 μl of each sample was spotted on nitrocellulose membrane, blocked with 2.5 % BSA, followed by immunoblotting analysis with according antibody.

### *In vitro* dephosphorylation assay

Tyrosine phosphorylated THEMIS protein was immunoprecipitated with FLAG resin from HEK293T cell co-expressing both FLAG-THEMIS and LCK. Recombinant SHP1-WT or SHP1-C453S proteins were added into immunoprecipitate above at the amount of 0, 0.25, 0.5, 1 and 2 μg, respectively. The reaction was kept at 30 °C water bath for 1 hour and stopped by adding 2x SDS loading buffer. Sample were heated at 95 °C for 5 mins and subjected to immunoblotting analysis.

### Mice

To establish THEMIS^-/-^ and THEMIS^Y34F/Y34F^ mice, we first generated Cas9 mRNA and sgRNA (Table 4). For *in vitro* transcription (IVT) of Cas9 mRNA and sgRNA, T7 promoter was added to the Cas9 coding region and sgRNA-scaffold by PCR amplification of plasmid PX330 (Addgene, #42230). PCR products of Cas9 and sgRNA region were used for IVT by following the manufactural of mMESSAGE mMACHINE T7 ULTRA kit (Life Technologies) and MEGA shortscript T7 kit (Life Technologies), respectively. Both Cas9 mRNA and sgRNAs were purified using the MEGA clear kit (Life Technologies) and stored at −80 °C for further experiments.

**Table 4:**
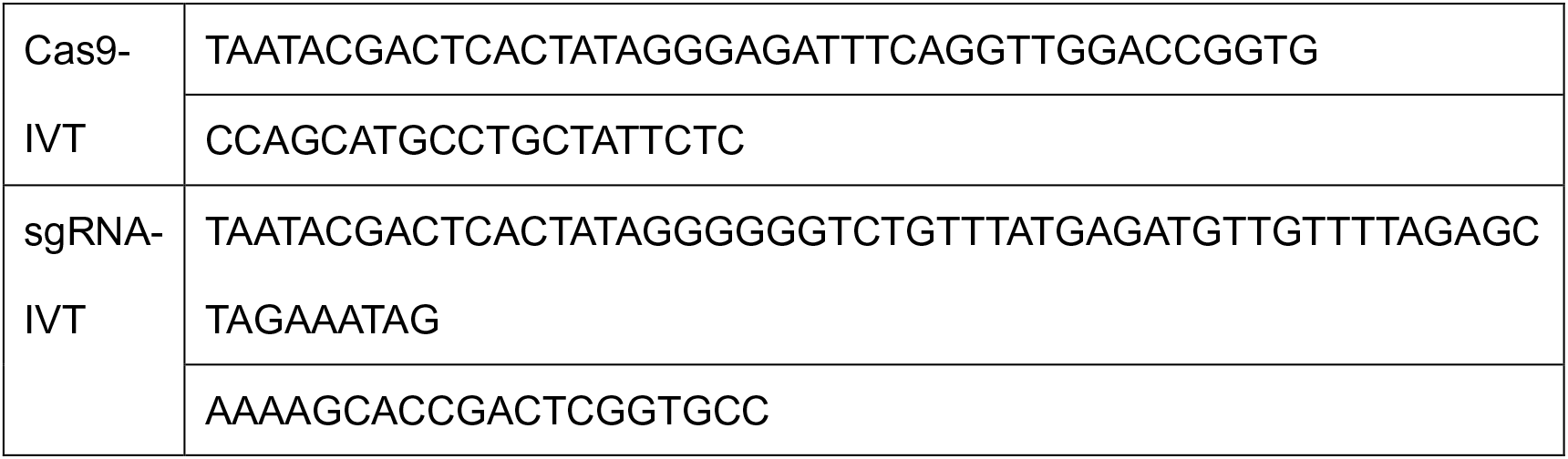
Cas9 mRNA and sgRNA for transgenic mice

For mice gene editing, superovulated female C57BL/6 mice were mated to C57BL/6 mice, fertilized embryos were collected from oviducts. Cas9 mRNA (100ng/µl), SgRNAs (50ng/µl for each sgRNA) and Oligos (100ng/µl) were mixed and injected into the cytoplasm of fertilized eggs in a droplet of HEPES-CZB medium containing 5ug/ml cytochalasin B (CB) using a FemtoJet microinjector (Eppendorf) with constant flow settings. The injected zygotes were cultured in KSOM medium with amino acids at 37°C, 5% CO_2_ until the 2-cell stage by 1.5 days. Thereafter, 25-30 2-cells-stage embryos were transferred into oviducts of pseudo-pregnant ICR females at 0.5 dpc. The tail ends of offspring were collected after born for DNA extraction and genotype analysis. The mice genotype was detected by Nest PCR and the primers were showed in Table 5.

**Table 5:**
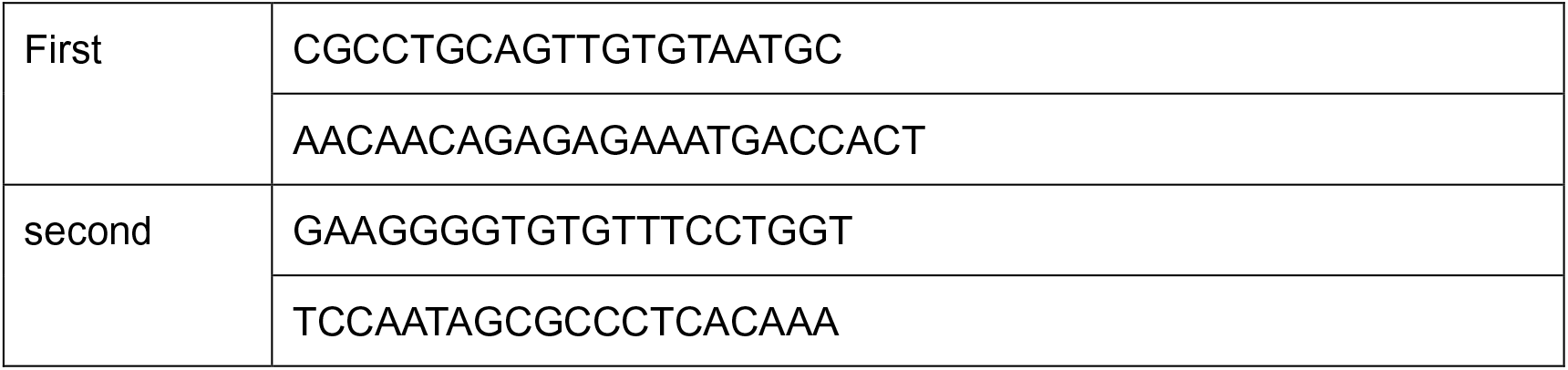
Primers of genotyping Nest PCR

We crossed THEMIS^-/-^ and THEMIS^Y34F/Y34F^ mice with OT1 mice to generate THEMIS^-/-^ OT1 and THEMIS^Y34F/Y34F^ OT1 mice, respectively. The transgenic T cell receptor of OT1 mice was designed to recognize ovalbumin peptide residues 257 to 264 (OVA257-264) in the context of H-2Kb (CD8 coreceptor interaction with MHC class I). This results in MHC class I restricted and ovalbumin-specific CD8+ T cells. The mice genotype was detected by PCR according to the protocol (Protocol 22849: Standard PCR Assay – Tg (Tcra)1100Mjb, The Jackson Laboratory).

All mice were raised in SPF (Specific Pathogen Free) condition at the National Facility for Protein Science of China in Shanghai (NFPS) Zhangjiang Lab. T cell development assays were performed using 6∼7-week old male and female mice. T cell homeostasis assays were performed using 12∼14-week old male and female mice.

### PEPSI (PUP-IT Enhanced Phosphatase Substrate Identification) assay

To generate inducible Pup (iPup) cell lines, we first produced lentivirus with Pup(E) plasmid Biotin-Pup(E)-IRES-BFP within the Tet-On 3G inducible expression system (Clontech 631168), and infected Jurkat cells for 48 hours. Then doxycycline (final concentration 2 μg/ml Selleck, S4136) was added into the culture medium for 24 hours. BFP-positive cells were sorted into 96-well plates by flow cytometry for single clone selection. It takes ∼3 weeks for cell re-population. After adding doxycycline (final concentration 2 μg/ml) and biotin (final concentration 4 μM) for 24 hours, the BFP expression of each clone was confirmed by flow cytometry, and the expression and modification of Bio-Pup(E) in BFP-positive cells was also confirmed by immunoblotting analysis.

We further subcloned SHP1-WT or SHP1-D419A into the PafA-IRES-puro-EGFP plasmid, respectively, and produced lentivirus to infect iPup Jurkat cells for 48 hrs. Cells were placed under puromycin selection (final concentration 2 μg/ml). The expression of SHP1-WT-PafA-Myc or SHP1-DA-PafA-Myc was confirmed by immunoblotting analysis.

SHP1-WT-PafA-Myc or SHP1-DA-PafA-Myc expressed iPup Jurkat cells were then grown in 75 cm^2^ flask. We added doxycycline (final concentration 2 μg/ml) and biotin (final concentration 4 μM) to the medium in advance, and induce expression for 24 hrs. Cells were lysed (50 mM Tris7.5, 200 mM NaCl, 2 % Triton, 0.1 % SDS and plus proteasome inhibitor cocktail), and biotin-pupE labelled proteins were enriched by streptavidin magnetic beads (NEB, S1420S) and identified by mass spectrometry analysis (Liu *et al*., 2018).

Particularly in this study, we compared the binding protein differences between SHP1-WT and SHP1-D419A in Jurkat cells. To obtain reliable and quantitative measurement, each group of samples was triplicated. To analyze the different proteins bound to SHP1-WT or SHP1-DA in Jurkat cells, the raw data were analyzed according to the procedure of the Perseus software under category “Unlabeled User Case”(Tyanova et al., 2016). We used GraphPad Prism to draw the relevant volcano maps. Same procedure was applied to both PTP1B-WT vs. PTP1B-DA and SHP2-DA vs. SHP1-DA PEPSI assays.

### Sample preparation for mass spectrometry

For identification of interacting proteins by mass spectrometry, immunoprecipitates from PEPSI assays were harvested, followed by on-beads trypsin digestion. In brief, before immunoprecipitation, lysates were added urea to a final concentration of 8M and treated with 10 mM (final concentration) DTT for 1 hr at 56°C and 25 mM IAM for 45 mins in dark to complete reductive alkylation. And then, lysates were also treated with 25 mM DTT 0.5 hour for further reduction. Adding 50 μl magnetic streptavidin beads (NEB, S1420S) for 1 hour to binding with the pupE labelled protein. Then, beads were collected by magnetic frame, with 2x sequential washes by Buffer 1-4 (Buffer 1: 8 M Urea, 50 mM Tris 8.0, 200 mM NaCl, 0.2% SDS; Buffer 2: 8 M Urea, 50 mM Tris 8.0, 200 mM NaCl; Buffer 3: 50 mM Tris 8.0, 0.5 mM EDTA, 1 mM DTT; Buffer 4: 50 mM NH_4_HCO_3_). Proteins were digested on beads with trypsin overnight at 37°C. Peptides were desalted by Ziptip, then vacuum-dried and re-suspended for following mass spectrometry characterization.

Mass spectrometry analysis was performed at the Proteomics Facility in Shanghaitech University. An Easy-nLC 1000 system coupled to a Q Exactive HF (both from Thermo Scientific) was used to separate and analyze peptides. The raw data were processed and searched with MaxQuant 1.5.4.1 with MS tolerance of 4.5 ppm, and MS/MS tolerance of 20 ppm. The UniProt human protein database (release 2016_07, 70630 sequences) and database for proteomics contaminants from MaxQuant were used for database search.

### Isolation of thymocyte and splenocytes by flow cytometry

Thymus and spleen were isolated from 6∼7-week old mice. To obtain single cell resuspension, both tissues were mashed, and strained with a 40 µm cell strainer into 5ml RPMI medium. Use Beckman-counter to count cell number, and take 8 million cells for following cell surface staining. Antibodies were diluted in cell staining buffer (4A Biotech, FXP005). Thymus (Qdot-605 CD4; APC-Cy7 CD8; APC CD44; BV421 CD25; PerCP-Cy5.5 TCRb; PE-Cy7 CD69; Zombie Violet); Spleen (PerCP-Cy5.5 CD4; FITC CD8; PE CD25; APC CD44; BV650 CD62L; Zombie UV). The staining procedure was performed at 4 °C for 1 hr. Cells were sorted by flow cytometry (BD Fortessa) and analyzed by FlowJo.

### Cell proliferation responds to IL7 & IL12 stimulation

Spleen and lymph nodes were isolated from mice and single cell resuspensions were obtained by mashing and straining with a 40 µm cell strainer. CD8^+^CD44^lo^ naïve T cells were sorted using immunomagnetic negative selection system (Stem cell, 19858), followed by CFSE (Beyotime, C1031) staining. Cells were resuspended into complete RPMI (cRPMI) medium which containing 10% FBS, 1% penicillin and streptomycin, 292 mg/ml l-glutamine, 50 μM 2-mercaptoehthanol (Gibco, 21985023), 1 mM sodium pyruvate (Gibco, 11360070), 1× MEM non-essential amino acids (100×, Gibco, 11140050) and 25 mM HEPES (Gibco, 25-0691-82). Then, 0.1 million cells were plated into 96-well U-bottom plate with 50 ng/ml IL7 (PeproTech, 217-17) and 50 ng/ml IL12 (PeproTech, 210-12)(Brzostek *et al*., 2020). The samples were cultured at 37 °C in an atmosphere of 5% CO_2_ for 7 days. Then, cells were staining with CD8 (mouse CD8-APC-Cy7, eBioscience, 47-0081-82) and Zombie UV (Biolegend, 427107), followed by flow cytometry analysis.

### T cell activation

Jurkat cells were first rested in RPMI medium without FBS for 1 hour, and then activated by anti-TCR antibody (C305) at different time points and stopped by double concentration of IP buffer (mentioned above).

For mouse T cells activation: cells expansion *in vitro* according to the procedure above in mouse T cell culture. Cells were rested in sDMEM medium without IL-2 cytokine for 24 hrs. For TCR activation, cells were incubated at 37 °C with 10 ug/ml anti-mCD3 (BioXcell, BE0001-1) 30s followed by 20 ug/ml AffiniPure Goat Anti-Armenian Hamster IgG (Jackson, 127-005-099) to stimulate cells at different time points and stopped by double concentration of cell lysis buffer (mentioned above). For IL7 and IL12 cytokines downstream activation, the samples were incubated at 37 °C with 50 ng/ml IL7 and IL12. Then, cell lysates were prepared for next immunoprecipitation or immunoblotting assays.

### Quantification and statistical analysis

Quantitative data were presented as mean ± SD. Statistical significance was assessed on the basis of p-values calculated via two-tailed unpaired Student’s t-test. Number of experiments (n) used for statistical evaluation was specified in relevant figure legends.

### Antibodies

For immunoprecipitation and immunoblotting analysis: anti-mouse THEMIS (Millipore, 06-1328), anti-human THEMIS (Sigma, HPA031425 and BD, 562587), anti-pY34 THEMIS (Generated by Abclonal, Wuhan, P.R. China; antigen: modified peptide EGSIpYEMFGNEC conjugated with KLH), anti-pTyr (4G10, Millipore, 05-321), anti-FLAG (GNI, GNI4110-FG), anti-Myc (Cell Signaling Technology, 2276), anti-LCK (Santa Cruz, SC-433), anti-pY416 SRC (Thermo Fisher, 44-660G and Cell Signaling Technology, 2101), anti-ERK1/2 (Cell Signaling Technology, 4377), anti-phospho-ERK1/2 (Thr202/Tyr204, Cell Signaling Technology, 92711), anti-SHP1 (Santa Cruz, SC-287; Cell Signaling Technology, 3759 and Santa Cruz, SC-7289), anti-SHP2 (Cell Signaling Technology, 3397), anti-ZAP70 (Santa Cruz, SC-32760), anti-AKT (Cell Signaling Technology, 4685), anti-pS473 AKT (Cell Signaling Technology, 4060), anti-Streptavidin-HRP (Cell Signaling Technology, 3999), anti-PTP1B (FG6), anti-Tubulin (Sigma, T6557), anti-Actin (Sigma, A2066) and anti-GAPDH-HRP (Abcam, ab105428).

For flow cytometry: anti-mouse CD4-Qdot-605 (Invitrogen, Q10092), anti-mouse CD8-APC-Cy7 (eBioscience, 47-0081-82), anti-mouse CD44-APC (Biolegend, 103012), anti-mouse CD25-BV421 (Biolegend, 102033), anti-mouse TCRb-PerCP-Cy5.5 (eBioscience, 45-5961-82), anti-mouse CD69-PE-Cy7 (eBioscience, 25-0691-82), anti-mouse CD4-PerCP-Cy5.5 (Biolegend, 116012), anti-mouse CD8-FITC (Biolegend, 100706), anti-mouse CD25-PE (eBioscience, 12-0051-83), anti-mouse CD62L-BV650 (BD, 564108), anti-mouse CD5-PE (eBioscience, 12-0051-83)

## Data availability

The mass spectrometry proteomics data have been deposited to the ProteomeXchange Consortium (http://proteomecentral.proteomexchange.org) via the iProX partner repository(Ma et al., 2019) with the dataset identifier PXD028689. Other raw and analyzed data files are available from the corresponding author upon reasonable request.

## Acknowledgments

The authors thank the staff members of the Animal Facility at the National Facility for Protein Science in Shanghai (NFPS), Zhangjiang Lab, China, for providing the support in mouse housing and care. The authors thank the Multi-Omics Core Facility (MOCF) at the School of Life Science and Technology, ShanghaiTech University for technical supports with mass spectrometry experiments. The authors also thank the Molecular and Cell Biology Core Facility (MCBCF) at the School of Life Science and Technology, ShanghaiTech University for technical support with flow cytometry. This work was supported by the Ministry of Science and Technology of China (2018YFC1004603 and SQ2021YFA080048 to GF), the National Natural Science Foundation of China (32070776 and 31872831 to GF, 31670919 to HW), the Shanghai Science and Technology Commission (19JC1413800 to GF), the Shanghai Shuguang program (19SG55 to GF) and a ShanghaiTech University startup grant.

## Author contribution

JZ, HW and GF designed the study. JZ, MS, LC, ZJ, SC, XX and YW performed the experiments, JZ was the major contributor; EZ generated THEMIS^Y34F/Y34F^ and THEMIS^-/-^ mice; PH, TH, MZ, LZ, HW and GF interpreted the data; JZ and GF wrote the paper with the help from HW.

The authors declare that they have no competing interests.

---

**Supplemental figure 1-related to Figure 1:**
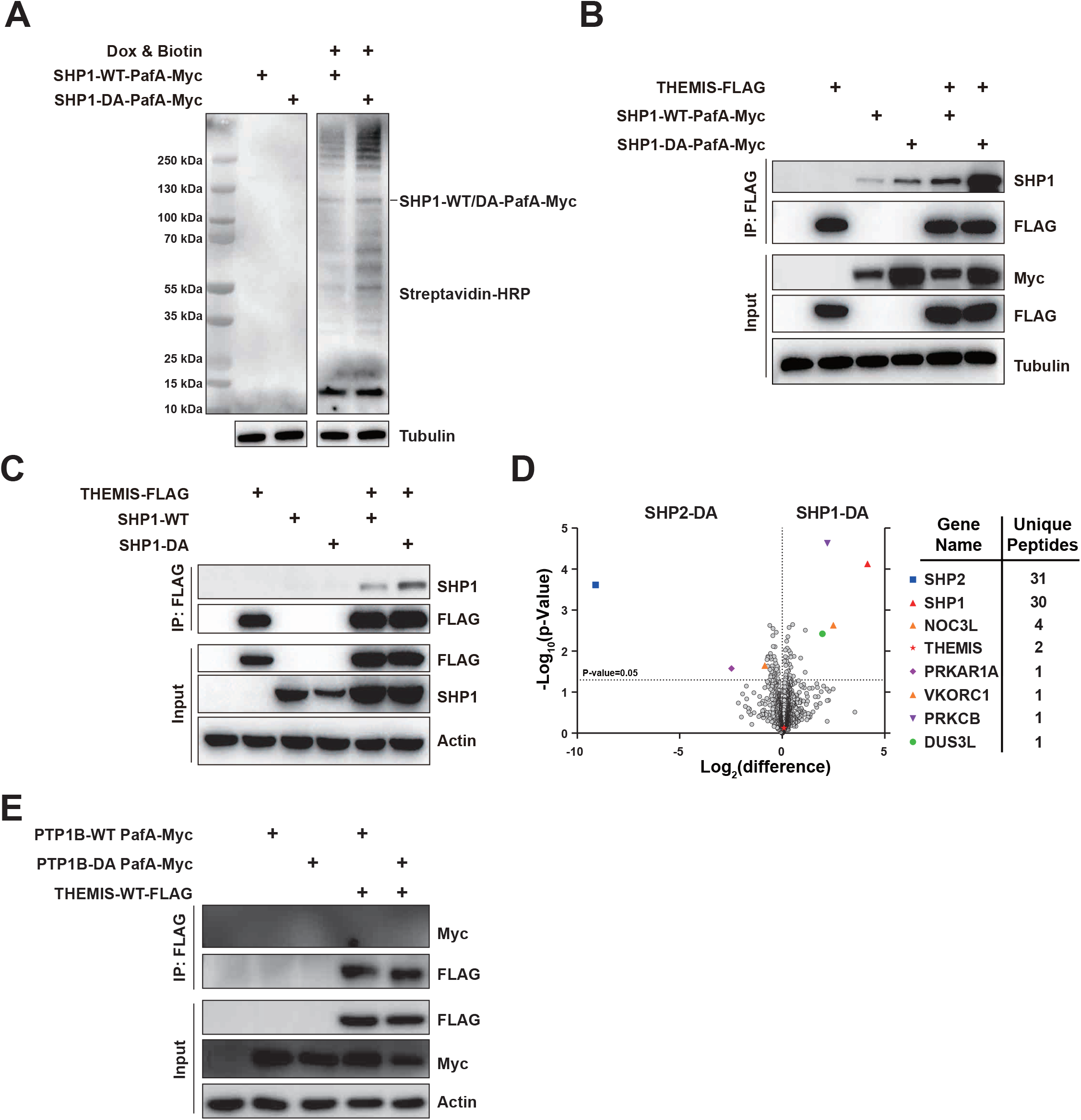
PEPSI strategy identified THEMIS as a novel substrate of non-receptor tyrosine phosphatase SHP1. (A) Inducible expressing pupE Jurkat T cells were stable transfected with SHP1-WT-PafA-Myc or SHP1-DA-PafA-Myc, indicated that PafA could successfully label SHP1 and SHP1 interactome with pupE which recruit biotin stained by streptavidin antibody. (B-C) HEK293T cells were transiently transfected with THEMIS-WT-FLAG and SHP1-WT-PafA-Myc or SHP1-DA-PafA-Myc (B) and SHP1-WT or SHP1-DA (C), indicated that THEMIS bound more with SHP1-DA than SHP1-WT. (D) A volcano plot of the SHP2-DA versus SHP1-DA PEPSI assay; the X axis represents the Log2 (difference) and the Y axis represents –Log10 (p-Value). The data was analyzed by Perseus_1.5.5.3 proteomics analysis tools. The significant hits identified in PEPSI assay are labelled using different colors and shapes, and ranked based on the number of unique hit peptides (E) HEK293T cells were transiently transfected with THEMIS-WT-FLAG and PTP1B-WT-PafA-Myc or PTP1B-DA-PafA-Myc. Results from co-immunoprecipitation assay indicated that PTP1B, either WT or DA mutant, couldn’t bind to THEMIS.

**Supplemental figure 2-related to Figure 2:**
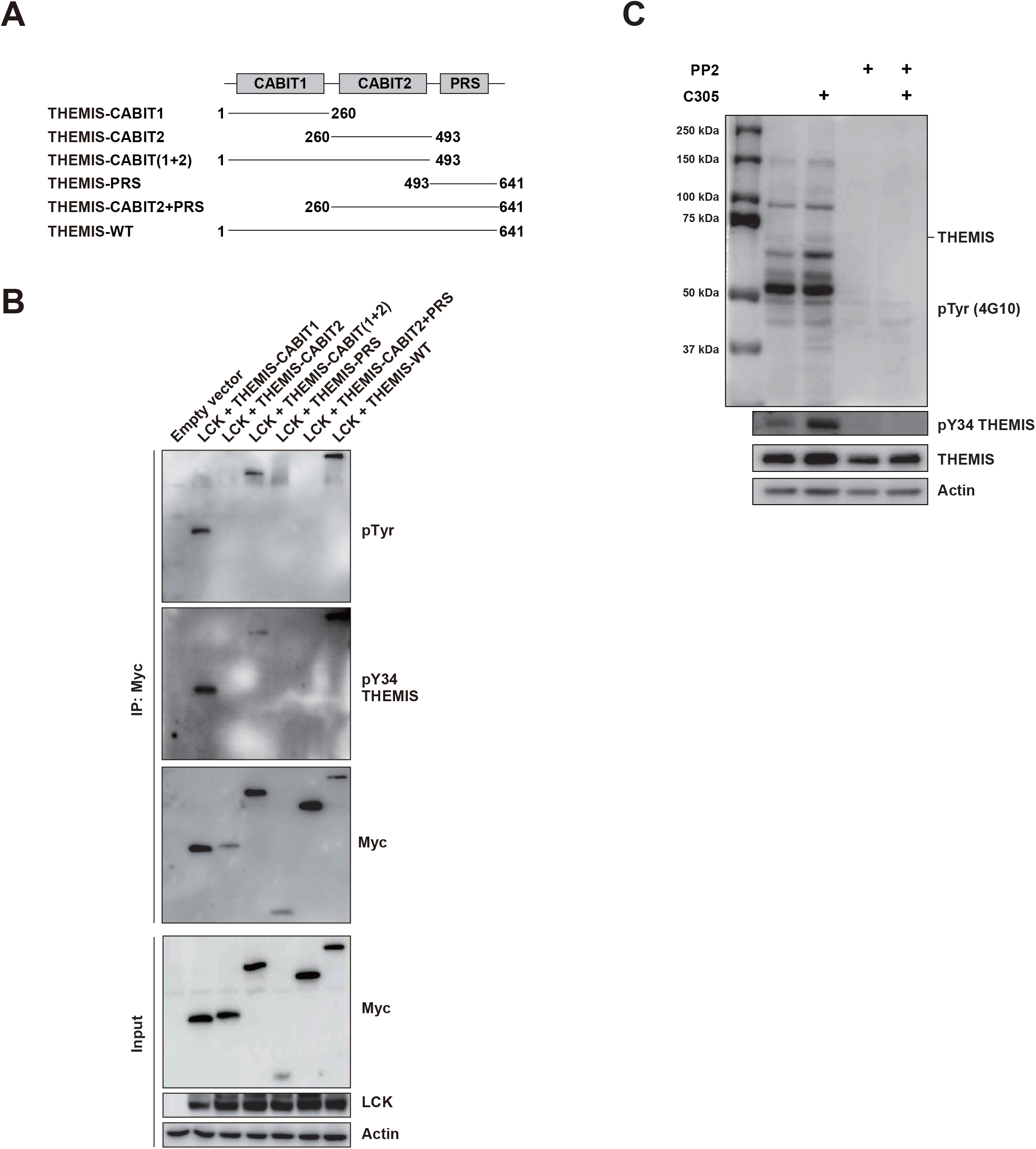
Phosphorylation regulation of THEMIS at Tyr34 site. (A-B) Immunoblotting of truncation variant experiments in the HEK293T cell line exploring the three domains of THEMIS with five different truncation mutants (A), Western blotting results showing that LCK can only phosphorylate the THEMIS Tyr34 site, positioned in the CABIT1 domain (B). (C) Src kinase family inhibitor PP2 treatment contributed to the blockade of phosphorylation transduction including THEMIS Tyr34 site after TCR signaling activation by C305 (anti-TCR) antibody.

**Supplemental figure 3-related to Figure 4:**
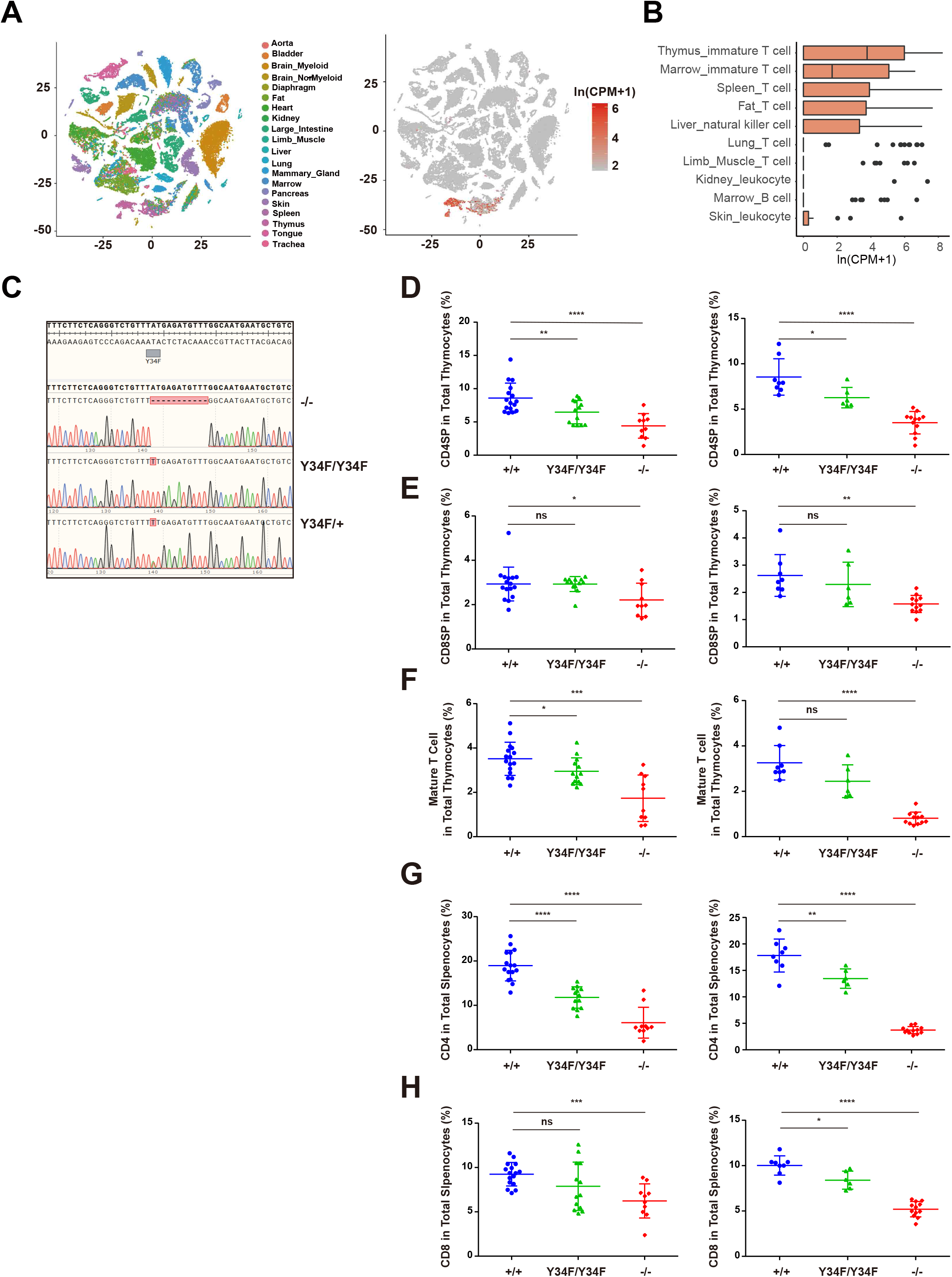
THEMIS^Y34F/Y34F^ mice showed a significant developmental defect in CD4 thymocyte development. (A) Distribution of t-SNE of single cell RNA-seq data from Tabula Muris, colored by different tissue (left) and expression of THEMIS among different tissues (right). (B) THEMIS expression in top 10 cell type ordered by the mean of THEMIS log transform expression in individual cell type, data from Tabula Muris. (C) Sanger sequencing of THEMIS^-/-^, THEMIS^Y34F/Y34F^ and THEMIS^Y34F/+^ mice tail genotype identification. (D-H) Mouse T cells (6∼7-week old) were stained with anti-CD4-Qdot605, anti-CD8-APC-Cy7, anti-TCRb-PerCP-Cy5.5, and anti-CD69-PE-Cy7 in thymocytes or CD4-PerCP-Cy5.5 and CD8-FITC in splenocytes for 1 hour on ice. Quantification of percentage of CD4SP (D), CD8SP (E), and mature cells (F) in total thymocytes or CD4+ (G) and CD8+ (H) T cells among total splenocytes isolated from male mice (left) and female mice (right) of the three genotypes (male: n=16, 13, 10; female: n =8, 6, 12). * P<0.05, ** P<0.01, *** P<0.001, **** P<0.0001 (two tail unpaired t-test)

**Supplemental figure 4-related to Figure 5:**
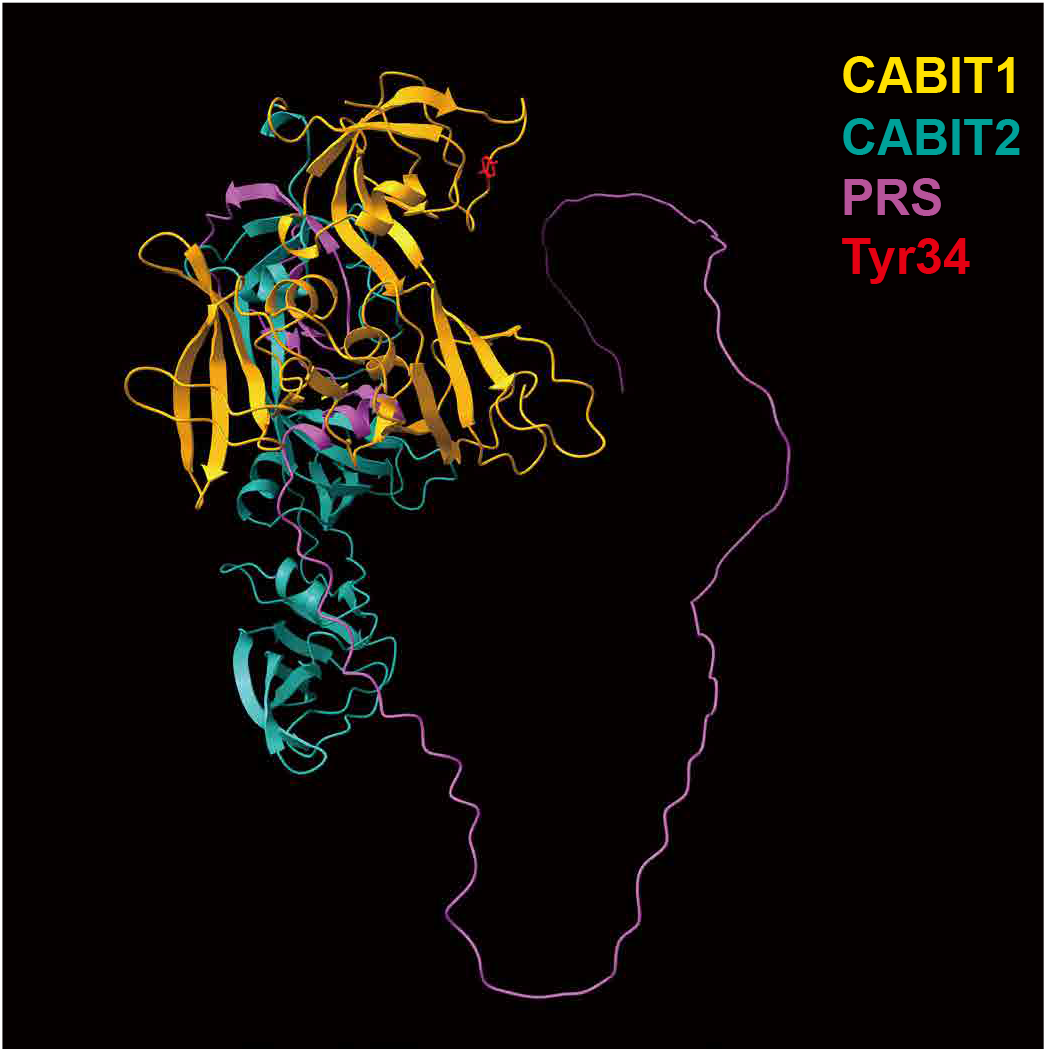
The structure of THEMIS predicted by AlphaFold, with Tyr34 highlighted. The CABIT1, CABIT2, PRS domains and Tyr34 residue were colored in orange, cyan, pink and red, respectively (https://www.alphafold.ebi.ac.uk/entry/Q8N1K5). Visualization of the atomic model was processed by UCSF ChimeraX.

## Notes

### Competing Interest Statement

The authors have declared no competing interest.

## References

1. Abram, C.L., and Lowell, C.A. (2017). Shp1 function in myeloid cells. J Leukoc Biol 102, 657–675. 10.1189/jlb.2MR0317-105R.

2. Allen, P.M. (2009). Themis imposes new law and order on positive selection. Nat Immunol 10, 805–806. 10.1038/ni0809-805.

3. Brugnera, E., Bhandoola, A., Cibotti, R., Yu, Q., Guinter, T.I., Yamashita, Y., Sharrow, S.O., and Singer, A. (2000). Coreceptor reversal in the thymus: signaled CD4+8+ thymocytes initially terminate CD8 transcription even when differentiating into CD8+ T cells. Immunity 13, 59–71. 10.1016/s1074-7613(00)00008-x.

4. Brzostek, J., Gautam, N., Zhao, X., Chen, E.W., Mehta, M., Tung, D.W.H., Chua, Y.L., Yap, J., Cho, S.H., Sankaran, S., et al. (2020). T cell receptor and cytokine signal integration in CD8(+) T cells is mediated by the protein Themis. Nat Immunol 21, 186–198. 10.1038/s41590-019-0570-3.

5. Choi, S., Cornall, R., Lesourne, R., and Love, P.E. (2017a). THEMIS: Two Models, Different Thresholds. Trends Immunol 38, 622–632. 10.1016/j.it.2017.06.006.

6. Choi, S., Warzecha, C., Zvezdova, E., Lee, J., Argenty, J., Lesourne, R., Aravind, L., and Love, P.E. (2017b). THEMIS enhances TCR signaling and enables positive selection by selective inhibition of the phosphatase SHP-1. Nat Immunol 18, 433–441. 10.1038/ni.3692.

7. Flint, A.J., Tiganis, T., Barford, D., and Tonks, N.K. (1997). Development of “substrate-trapping” mutants to identify physiological substrates of protein tyrosine phosphatases. Proc Natl Acad Sci U S A 94, 1680–1685. 10.1073/pnas.94.5.1680.

8. Fry, T.J., Moniuszko, M., Creekmore, S., Donohue, S.J., Douek, D.C., Giardina, S., Hecht, T.T., Hill, B.J., Komschlies, K., Tomaszewski, J., et al. (2003). IL-7 therapy dramatically alters peripheral T-cell homeostasis in normal and SIV-infected nonhuman primates. Blood 101, 2294–2299. 10.1182/blood-2002-07-2297.

9. Fu, G., Casas, J., Rigaud, S., Rybakin, V., Lambolez, F., Brzostek, J., Hoerter, J.A., Paster, W., Acuto, O., Cheroutre, H., et al. (2013). Themis sets the signal threshold for positive and negative selection in T-cell development. Nature 504, 441–445. 10.1038/nature12718.

10. Fu, G., Vallee, S., Rybakin, V., McGuire, M.V., Ampudia, J., Brockmeyer, C., Salek, M., Fallen, P.R., Hoerter, J.A., Munshi, A., et al. (2009). Themis controls thymocyte selection through regulation of T cell antigen receptor-mediated signaling. Nat Immunol 10, 848–856. 10.1038/ni.1766.

11. Gascoigne, N.R., and Acuto, O. (2015). THEMIS: a critical TCR signal regulator for ligand discrimination. Curr Opin Immunol 33, 86–92. 10.1016/j.coi.2015.01.020.

12. Gascoigne, N.R., Brzostek, J., Mehta, M., and Acuto, O. (2016a). SHP1-ing thymic selection. Eur J Immunol 46, 2091–2094. 10.1002/eji.201646582.

13. Gascoigne, N.R., Rybakin, V., Acuto, O., and Brzostek, J. (2016b). TCR Signal Strength and T Cell Development. Annu Rev Cell Dev Biol 32, 327–348. 10.1146/annurev-cellbio-111315-125324.

14. Germain, R.N. (2002). T-cell development and the CD4-CD8 lineage decision. Nat Rev Immunol 2, 309–322. 10.1038/nri798.

15. Godfrey, D.I., Kennedy, J., Suda, T., and Zlotnik, A. (1993). A developmental pathway involving four phenotypically and functionally distinct subsets of CD3-CD4-CD8-triple-negative adult mouse thymocytes defined by CD44 and CD25 expression. J Immunol 150, 4244–4252.

16. Goplen, N.P., Saxena, V., Knudson, K.M., Schrum, A.G., Gil, D., Daniels, M.A., Zamoyska, R., and Teixeiro, E. (2016). IL-12 Signals through the TCR To Support CD8 Innate Immune Responses. J Immunol 197, 2434–2443. 10.4049/jimmunol.1600037.

17. Hsu, P.D., Lander, E.S., and Zhang, F. (2014). Development and applications of CRISPR-Cas9 for genome engineering. Cell 157, 1262–1278. 10.1016/j.cell.2014.05.010.

18. Jinek, M., Chylinski, K., Fonfara, I., Hauer, M., Doudna, J.A., and Charpentier, E. (2012). A programmable dual-RNA-guided DNA endonuclease in adaptive bacterial immunity. Science 337, 816–821. 10.1126/science.1225829.

19. Johnson, A.L., Aravind, L., Shulzhenko, N., Morgun, A., Choi, S.Y., Crockford, T.L., Lambe, T., Domaschenz, H., Kucharska, E.M., Zheng, L., et al. (2009). Themis is a member of a new metazoan gene family and is required for the completion of thymocyte positive selection. Nat Immunol 10, 831–839. 10.1038/ni.1769.

20. Kakugawa, K., Yasuda, T., Miura, I., Kobayashi, A., Fukiage, H., Satoh, R., Matsuda, M., Koseki, H., Wakana, S., Kawamoto, H., and Yoshida, H. (2009). A novel gene essential for the development of single positive thymocytes. Mol Cell Biol 29, 5128–5135. 10.1128/MCB.00793-09.

21. Lesourne, R., Uehara, S., Lee, J., Song, K.D., Li, L., Pinkhasov, J., Zhang, Y., Weng, N.P., Wildt, K.F., Wang, L., et al. (2009). Themis, a T cell-specific protein important for late thymocyte development. Nat Immunol 10, 840–847. 10.1038/ni.1768.

22. Liu, Q., Zheng, J., Sun, W., Huo, Y., Zhang, L., Hao, P., Wang, H., and Zhuang, M. (2018). A proximity-tagging system to identify membrane protein-protein interactions. Nat Methods 15, 715–722. 10.1038/s41592-018-0100-5.

23. Ma, J., Chen, T., Wu, S., Yang, C., Bai, M., Shu, K., Li, K., Zhang, G., Jin, Z., He, F., et al. (2019). iProX: an integrated proteome resource. Nucleic Acids Res 47, D1211–D1217. 10.1093/nar/gky869.

24. Mehta, M., Brzostek, J., Chen, E.W., Tung, D.W.H., Chen, S., Sankaran, S., Yap, J., Rybakin, V., and Gascoigne, N.R.J. (2018). Themis-associated phosphatase activity controls signaling in T cell development. Proc Natl Acad Sci U S A 115, E11331–E11340. 10.1073/pnas.1720209115.

25. Montalibet, J., Skorey, K.I., and Kennedy, B.P. (2005). Protein tyrosine phosphatase: enzymatic assays. Methods 35, 2–8. 10.1016/j.ymeth.2004.07.002.

26. Neel, B.G., Gu, H., and Pao, L. (2003). The ‘Shp’ing news: SH2 domain-containing tyrosine phosphatases in cell signaling. Trends Biochem Sci 28, 284–293. 10.1016/S0968-0004(03)00091-4.

27. O’Reilly, A.M., Pluskey, S., Shoelson, S.E., and Neel, B.G. (2000). Activated mutants of SHP-2 preferentially induce elongation of Xenopus animal caps. Mol Cell Biol 20, 299–311. 10.1128/MCB.20.1.299-311.2000.

28. Paster, W., Bruger, A.M., Katsch, K., Gregoire, C., Roncagalli, R., Fu, G., Gascoigne, N.R., Nika, K., Cohnen, A., Feller, S.M., et al. (2015). A THEMIS:SHP1 complex promotes T-cell survival. EMBO J 34, 393–409. 10.15252/embj.201387725.

29. Patrick, M.S., Oda, H., Hayakawa, K., Sato, Y., Eshima, K., Kirikae, T., Iemura, S., Shirai, M., Abe, T., Natsume, T., et al. (2009). Gasp, a Grb2-associating protein, is critical for positive selection of thymocytes. Proc Natl Acad Sci U S A 106, 16345–16350. 10.1073/pnas.0908593106.

30. Singer, A. (2002). New perspectives on a developmental dilemma: the kinetic signaling model and the importance of signal duration for the CD4/CD8 lineage decision. Curr Opin Immunol 14, 207–215. 10.1016/s0952-7915(02)00323-0.

31. Singer, A., Adoro, S., and Park, J.H. (2008). Lineage fate and intense debate: myths, models and mechanisms of CD4-versus CD8-lineage choice. Nat Rev Immunol 8, 788–801. 10.1038/nri2416.

32. Sprent, J., and Surh, C.D. (2011). Normal T cell homeostasis: the conversion of naive cells into memory-phenotype cells. Nat Immunol 12, 478–484. 10.1038/ni.2018.

33. Sun, C., Shou, P., Du, H., Hirabayashi, K., Chen, Y., Herring, L.E., Ahn, S., Xu, Y., Suzuki, K., Li, G., et al. (2020). THEMIS-SHP1 Recruitment by 4-1BB Tunes LCK-Mediated Priming of Chimeric Antigen Receptor-Redirected T Cells. Cancer Cell 37, 216–225 e216. 10.1016/j.ccell.2019.12.014.

34. Tabula Muris, C., Overall, c., Logistical, c., Organ, c., processing Library, p., sequencing, Computational data, a., Cell type, a., Writing, g., et al. (2018). Single-cell transcriptomics of 20 mouse organs creates a Tabula Muris. Nature 562, 367–372. 10.1038/s41586-018-0590-4.

35. Tamarit, B., Bugault, F., Pillet, A.H., Lavergne, V., Bochet, P., Garin, N., Schwarz, U., Theze, J., and Rose, T. (2013). Membrane microdomains and cytoskeleton organization shape and regulate the IL-7 receptor signalosome in human CD4 T-cells. J Biol Chem 288, 8691–8701. 10.1074/jbc.M113.449918.

36. Tyanova, S., Temu, T., Sinitcyn, P., Carlson, A., Hein, M.Y., Geiger, T., Mann, M., and Cox, J. (2016). The Perseus computational platform for comprehensive analysis of (prote)omics data. Nat Methods 13, 731–740. 10.1038/nmeth.3901.

37. Wang, W., Liu, L., Song, X., Mo, Y., Komma, C., Bellamy, H.D., Zhao, Z.J., and Zhou, G.W. (2011). Crystal structure of human protein tyrosine phosphatase SHP-1 in the open conformation. J Cell Biochem 112, 2062–2071. 10.1002/jcb.23125.

38. Welte, S., Baringhaus, K.H., Schmider, W., Muller, G., Petry, S., and Tennagels, N. (2005). 6,8-Difluoro-4-methylumbiliferyl phosphate: a fluorogenic substrate for protein tyrosine phosphatases. Anal Biochem 338, 32–38. 10.1016/j.ab.2004.11.047.

39. Yang, J., Liu, L., He, D., Song, X., Liang, X., Zhao, Z.J., and Zhou, G.W. (2003). Crystal structure of human protein-tyrosine phosphatase SHP-1. J Biol Chem 278, 6516–6520. 10.1074/jbc.M210430200.

40. Yu, Q., Erman, B., Bhandoola, A., Sharrow, S.O., and Singer, A. (2003). In vitro evidence that cytokine receptor signals are required for differentiation of double positive thymocytes into functionally mature CD8+ T cells. J Exp Med 197, 475–487. 10.1084/jem.20021765.

41. Zuniga-Pflucker, J.C. (2004). T-cell development made simple. Nat Rev Immunol 4, 67–72. 10.1038/nri1257.

